# The ubiquitin-conjugating enzyme Ubc13-Mms2 cooperates with a family of FYVE-type-RING ubiquitin protein ligases in K63-polyubiquitylation at internal membranes

**DOI:** 10.1101/575241

**Authors:** Christian Renz, Vera Tröster, Thomas K. Albert, Olivier Santt, Susan C. Jacobs, Anton Khmelinskii, Helle D. Ulrich

## Abstract

The heterodimeric ubiquitin-conjugating enzyme (E2), Ubc13-Mms2, catalyses K63-specific polyubiquitylation in genome maintenance and inflammatory signalling. In budding yeast, the only ubiquitin protein ligase (E3) known to cooperate with Ubc13-Mms2 so far is a nuclear RING finger protein, Rad5, involved in the replication of damaged DNA. We have now identified a family of membrane-associated FYVE-(type)-RING finger proteins as cognate E3s for Ubc13-Mms2 in several species. We show that budding yeast Pib1, a FYVE-RING finger E3 associated with internal membranes, exhibits exquisite selectivity for Ubc13-Mms2 and cooperates with the E2 in the multivesicular body pathway. Phenotypic analysis indicates that the contribution of Ubc13-Mms2 to membrane trafficking goes beyond its cooperation with Pib1, suggesting an involvement with additional E3s in the endocytic compartment. These results widely implicate Ubc13-Mms2 in the regulation of membrane protein sorting.

## Introduction

Posttranslational modification with ubiquitin serves as a versatile means to modulate protein function, not least because of the diversity of ubiquitin signals (Kwon & Ciechanover, 2017). Ubiquitin is usually attached to its substrates via an isopeptide bond between its carboxy (C)-terminus and an internal lysine (K) residue of the target protein. Modification is accomplished by the successive action of a ubiquitin-activating enzyme (E1), a ubiquitin-conjugating enzyme (E2) and a ubiquitin protein ligase (E3). Ubiquitin itself can serve as a target for further modification. Depending on which of its seven lysine residues is used as an acceptor for conjugation, polyubiquitin chains are formed with varying topology (Komander & Rape, 2012). Even ubiquitin’s amino (N)-terminus can be modified, resulting in a head-to-tail linkage (Rieser et al, 2013). Downstream effector proteins that recognise the modification via dedicated ubiquitin-binding domains mediate the biological effects (Husnjak & Dikic, 2012). As a consequence, ubiquitylation impinges on the conformation, localisation or interactions of the modified protein. As some ubiquitin-binding domains exhibit high selectivity towards defined linkages, the topology of a polyubiquitin chain is thought to play an important role in specifying the consequences of ubiquitylation. Whereas K48-, K11- and K29-linked polyubiquitylation has been implicated in proteasomal degradation, the K63-linkage is involved in the regulation of numerous non-proteasomal pathways ranging from the DNA damage response to inflammatory signalling, endocytosis, intracellular membrane trafficking and lysosomal targeting (Erpapazoglou et al, 2014; Garcia-Rodriguez et al, 2016; Panier & Durocher, 2009; Wu & Karin, 2015).

The topology of a polyubiquitin chain is determined by the enzymes involved in its assembly (Suryadinata et al, 2014). Among the characterised E2s, Ubc13 is unique in exclusively generating K63-linked chains. Its linkage specificity arises from an obligatory interaction partner, the catalytically inactive E2 variant Mms2 (Hofmann & Pickart, 1999). Ubc13 cooperates with a number of E3s, such as human TRAF6, CHFR, RNF8, RNF168 and CHIP (Hodge et al, 2016). In the budding yeast, *Saccharomyces cerevisiae*, the only E3 known to function with Ubc13 is Rad5, involved in the pathway of DNA damage bypass (Ulrich & Jentsch, 2000). All these E3s belong to the RING finger or the related U-box families and, in each case, Ubc13 apparently dictates linkage specificity (Zheng & Shabek, 2017). When combined with other E2s, some RING E3s such as CHIP can assemble polyubiquitin chains of alternative or even mixed linkages (Kim et al, 2007). In contrast to the RING and U-box proteins, E3s of the HECT and RBR families directly determine chain linkage. For example, the budding yeast HECT E3, Rsp5, predominantly catalyses ubiquitin polymerisation via K63, and the human RBR E3 LUBAC specifically assembles M1-linked chains (Erpapazoglou et al, 2014; Rieser et al, 2013). Most probably, the influence of these E3 families on linkage selectivity is due to their direct participation in the thioester-mediated transfer of ubiquitin to the acceptor lysine.

Together, Rsp5 (or its mammalian homologue NEDD4) and Ubc13 appear to be responsible for the bulk of K63-polyubiquitylation in yeast and vertebrate cells. Rsp5/NEDD4 is mainly involved in membrane trafficking, mediating ubiquitylation of various plasma membrane proteins and components of the endocytic machinery (Erpapazoglou et al, 2014). Human UBC13 cooperates with the E2 variant UBE2V2 (also called MMS2) in genome maintenance-related functions in the nucleus, while in the cytoplasm it associates with UBE2V1 (also called UEV1A) for inflammatory signalling in the NFκB pathway (Andersen et al, 2005). Surprisingly, in budding yeast, which lacks an NFκB pathway, Ubc13-Mms2 also localises mainly to the cytoplasm and accumulates in the nucleus only in response to DNA damage (Ulrich & Jentsch, 2000), thus raising the question of its cytoplasmic function.

We now report that Ubc13-Mms2 from *S. cerevisiae* acts as the cognate E2 for a poorly characterised RING E3, Pib1, which localises to endosomal and vacuolar membranes via a lipid-binding FYVE domain (Burd & Emr, 1998; Shin et al, 2001). Pib1 plays a role in the multivesicular body (MVB) pathway where it acts redundantly with an Rsp5 adaptor, Bsd2, in delivering plasma membrane proteins to the vacuole (Nikko & Pelham, 2009). We show that Pib1 is highly selective for Ubc13-Mms2 in terms of both physical interaction and activation of ubiquitin transfer, and that the proteins cooperate *in vivo* as a K63-specific E2-E3 pair. Moreover, we found a comparable activity with Ubc13-Mms2 in a series of other RING finger proteins associated with membranes of the endocytic and lysosomal compartments. Some of these share Pib1’s FYVE-RING domain arrangement, such as the postulated homologue from *Schizosaccharomyces pombe* (SpPib1) (Kampmeyer et al, 2017), an RNA-binding protein from *Ustilago maydis* (Upa1) (Pohlmann et al, 2015), and human Rififylin (also known as CARP2 or Sakura), which harbours a FYVE-like domain (Coumailleau et al, 2004). In combination with an analysis of genetic interactions, our data widely implicate Ubc13-Mms2 in the regulation of intracellular membrane protein sorting and turnover.

## Results

### Ubc13 interacts with the dimeric RING finger protein Pib1

In order to identify potential regulators of the heterodimeric E2 complex Ubc13-Mms2, we performed a two-hybrid screen using the *S. cerevisiae* E2 variant Mms2 as a bait. From a budding yeast genomic library of ~ 5·10^6^ clones (James et al, 1996) we obtained at least five independent isolates encoding the full sequence of Pib1. This 286 amino acid (aa) protein harbours an N-terminal FYVE domain and a C-terminal RING finger (Figure 1A) (Burd & Emr, 1998). When analysed individually in two-hybrid assays, Pib1 exhibited a moderate interaction with Mms2 in only one of the two orientations of the system, but interacted more strongly with Ubc13 (Figure 1B). Mutation of conserved Pib1 residues critical for E2 interaction in other RING finger proteins (C225A, I227A) (Metzger et al, 2014) abolished association with both Ubc13 and Mms2 (Figure S1A), suggesting an involvement of the RING finger in E2 binding.

**Figure 1.**
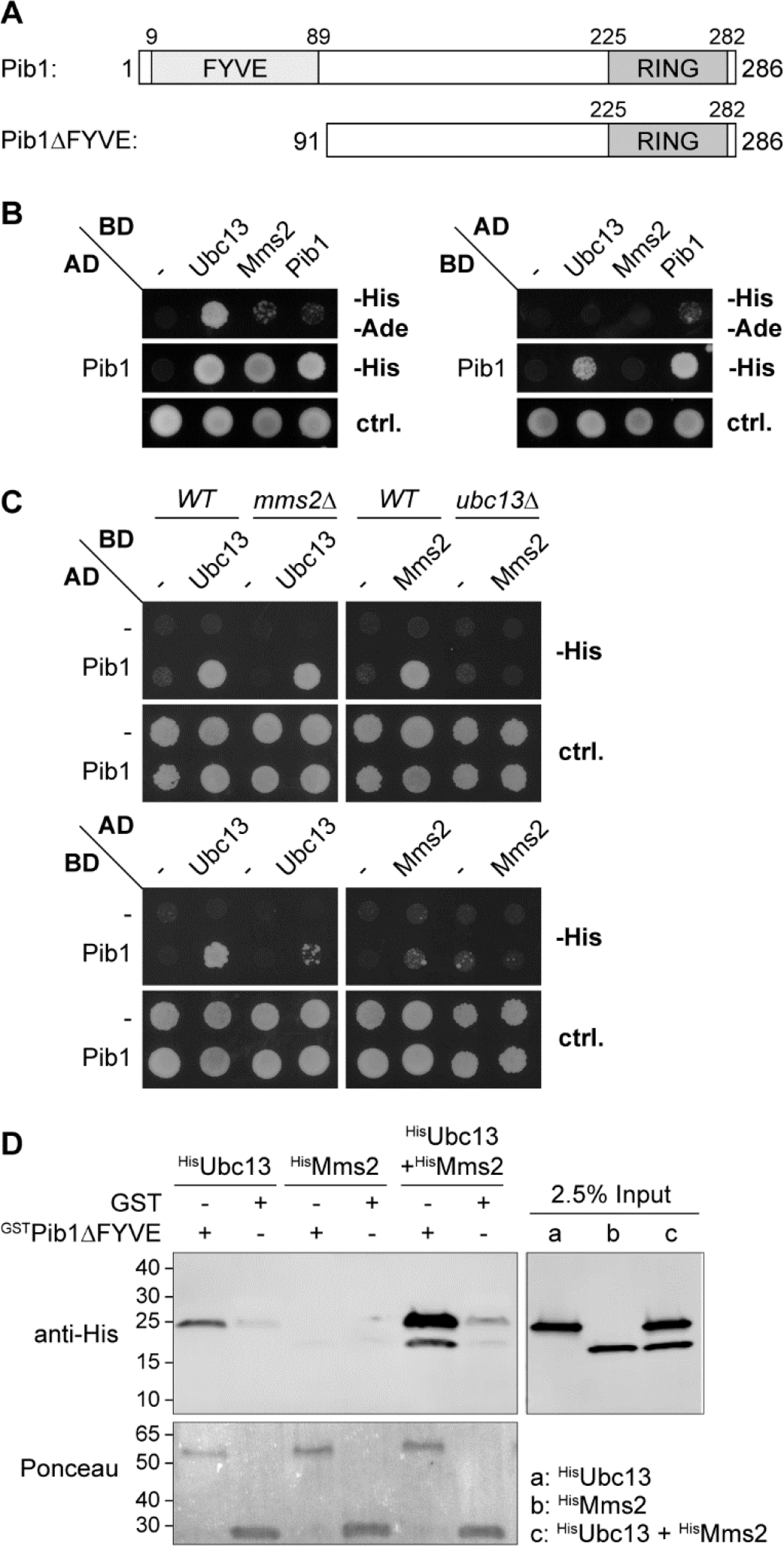
Pib1 interacts with Ubc13 in a manner that is enhanced by Mms2. **(A)** Schematic representation of Pib1 domain structure. **(B)** Interactions of Pib1 with Ubc13, Mms2 and itself in the two-hybrid assay (AD: Gal4 activation domain; BD: Gal4 DNA-binding domain). Growth on -His medium indicates positive interactions; growth on -His-Ade medium indicates strong associations. **(C)** Pib1-Mms2 interaction is mediated by Ubc13. Two-hybrid analysis was performed as in (B) in *mms2Δ* or *ubc13Δ* mutant reporter strains. **(D)** Mms2 requires Ubc13 for interaction with Pib1 *in vitro*. Association of recombinant ^GST^Pib1ΔFYVE with ^His^Ubc13 and ^His^Mms2 was assessed by pre-loading ^GST^Pib1ΔFYVE or GST onto glutathione Sepharose before incubation with 1 μM recombinant ^His^Ubc13, ^His^Mms2 or ^His^Ubc13-^His^Mms2 complex. Bound material was detected by western blotting with an anti-His_6_-tag antibody and Ponceau staining of the membrane.

The robust interaction between Pib1 and Ubc13 pointed to a direct association, whereas the weak interaction between Pib1 and Mms2 might reflect an indirect contact mediated by endogenous Ubc13 in the two-hybrid assay. We therefore deleted either *UBC13* or *MMS2* in the reporter strain (Figure 1C). Consistent with an Ubc13-mediated indirect interaction between Pib1 and Mms2, the two-hybrid signal was abolished in the *ubc13Δ* background. Surprisingly, deletion of *MMS2* also weakened the interaction between Pib1 and Ubc13 in one orientation, possibly suggesting a contribution of Mms2 to the interaction between Pib1 and the E2. Mutations in Ubc13 corresponding to positions that generally mediate RING binding in E2s, such as K6, K10, M64 and S96, also affected the Pib1-Ubc13 interaction (Figure S1B). This indicates that Ubc13 contacts the Pib1 RING domain in a “standard” orientation, i.e. comparable to its interaction with other E3s, including Rad5 (Metzger et al, 2014; Ulrich, 2003). We confirmed the direct interaction of Pib1 with Ubc13-Mms2 *in vitro* by immobilising recombinant, GST-tagged Pib1ΔFYVE on glutathione Sepharose and examining retention of the E2. Consistent with the two-hybrid data, Ubc13 alone or in complex with Mms2 bound to ^GST^Pib1ΔFYVE, whereas Mms2 failed to interact with ^GST^Pib1ΔFYVE in the absence of Ubc13 (Figure 1D). Again, the presence of Mms2 enhanced the interaction between Ubc13 and Pib1.

In the two-hybrid assay, association of Pib1 with itself was also noted (Figure 1B). Size exclusion chromatography analysis of a series of recombinant Pib1 truncation constructs revealed a running behaviour consistent with dimerisation, mediated by a sequence N-terminal to and including the RING finger. The minimal region adjacent to the RING domain required for robust dimerisation was found to be between 50 and 100 amino acids (Figure S1C).

### Pib1 is an E3 with high selectivity for Ubc13-Mms2

Our results suggested that Pib1 acts as a cognate E3 for Ubc13-Mms2. Indeed, *in vitro* ubiquitin polymerisation by Ubc13-Mms2 was strongly stimulated by the addition of Pib1ΔFYVE in a concentration-dependent manner (Figure 2A). Chain formation proceeded exclusively via K63, as mutant ubiquitin, Ub(K63R), was not polymerised at all, and the K63-specific ubiquitin isopeptidase AMSH (McCullough et al, 2004) efficiently disassembled the products of the Pib1-stimulated reaction (Figure 2B). Comparison of the *in vitro* activities of Pib1 truncation constructs showed a correlation between stimulation of polyubiquitin chain synthesis and Pib1 dimerisation, indicating that Pib1 is more effective as a dimer than as a monomer, but its FYVE domain is not required for catalytic activity (Figure S2A).

**Figure 2.**
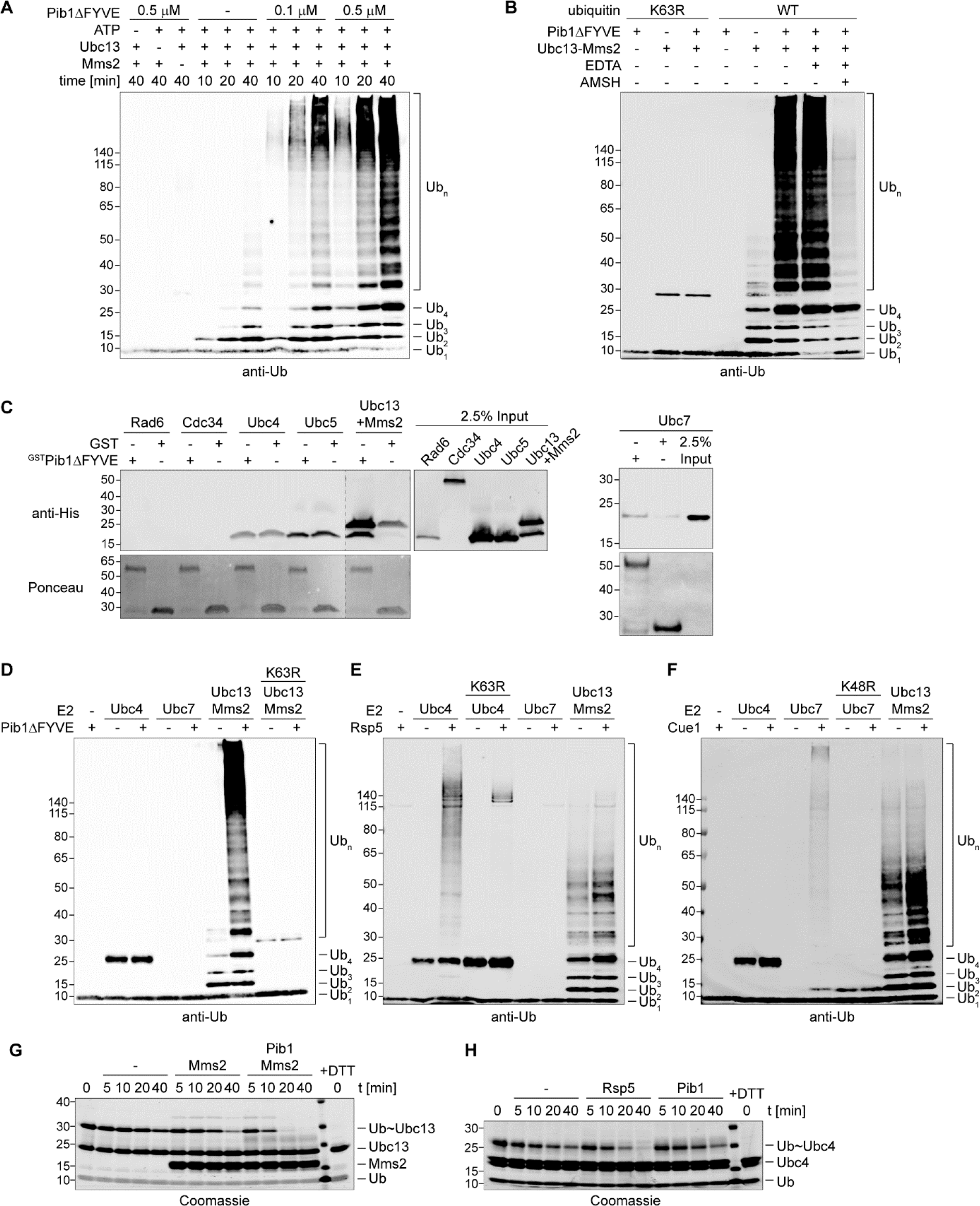
Pib1 is an E3 with high selectivity for Ubc13-Mms2. **(A)** Pib1 stimulates *in vitro* polyubiquitin chain synthesis by Ubc13-Mms2. Recombinant ^GST^Pib1ΔFYVE, ^His^Ubc13 and untagged Mms2 as well as ATP were added as indicated to reactions containing E1 and ubiquitin and incubated at 30°C for the indicated times. Products were analysed by western blotting with an anti-ubiquitin antibody. **(B)** Pib1 selectively stimulates the synthesis of K63-linked polyubiquitin chains. *In vitro* ubiquitylation reactions were set up as in (A), using WT or a K63R mutant ubiquitin, and incubated at 30°C for 30 min, followed by addition of 20 mM EDTA and further incubation in the presence or absence of the K63-specific ubiquitin isopeptidase ^GST^AMSH at 37°C for 60 min. **(C)** Pib1 selectively interacts with Ubc13-Mms2. Interaction of ^GST^Pib1ΔFYVE with ^His^Rad6, ^His^Cdc34, ^His^Ubc4, ^His^Ubc5, ^His^Ubc13-^His^Mms2 and ^His^Ubc7 was analysed as described in Figure 1D. **(D-F)** Pib1, Rsp5 and Cue1 exhibit distinct E2 and linkage selectivities in polyubiquitin chain synthesis. *In vitro* ubiquitylation assays were carried out at 30°C for 60 min as described in (A), using ^His^Ubc4, Ubc7^His^ and ^His^Ubc13-Mms2 as E2s and ^GST^Pib1ΔFYVE, ^GST^Rsp5 and ^GST^Cue1^His^ as E3s, respectively. Use of mutant ubiquitin is indicated by K63R or K48R, respectively. **(G)** Discharge of the Ubc13-thioester is enhanced by Pib1. Ubc13(K92R) was charged with ubiquitin in the presence of E1, ATP and ubiquitin for 60 min at 30°C. Reactions were stopped by apyrase treatment, and discharge of Ubc13~Ub was initiated by the addition of lysine, Mms2 and Pib1-RING+100aa as indicated. Reactions were incubated at 30°C for the indicated times. **(H)** Ubc4-thioester discharge is stimulated by Rsp5, but not by Pib1. Ubc4(K91R) was charged with ubiquitin for 10 min at 30°C, and discharge of Ubc4~Ub was analysed in the presence of lysine and Rsp5 or Pib1-RING+100aa as indicated.

As some E3s can function with multiple E2s, we asked how specific the Pib1-Ubc13-Mms2 interaction was. In fact, Pib1 had previously been reported to cooperate with one of the most non-selective human E2s, UBCH5A, *in vitro* (Shin et al, 2001); however, these observations were made with GST-tagged versions of both Pib1 and the E2, which might have induced inappropriate E2-E3 pairing. In a systematic two-hybrid assay with all budding yeast E2s, Pib1 exclusively interacted with Ubc13 and Mms2 (Figure S2B-D). *In vitro* interaction assays with ^GST^Pib1ΔFYVE showed no binding to a range of recombinant yeast E2s, including Rad6, Cdc34, Ubc7 and two highly promiscuous UBCH5A homologs, Ubc4 and Ubc5 (Figure 2C). Moreover, and consistent with a high selectivity for Ubc13-Mms2, Pib1ΔFYVE did not stimulate ubiquitin polymerisation by Ubc4 or Ubc7 (Figure 2D), even though both E2s were active in combination with appropriate E3s, Rsp5 and Cue1 (Bazirgan & Hampton, 2008), respectively (Figure 2E, F). Notably, those two E3s activated only their cognate E2, but not Ubc13-Mms2, thus indicating selectivity on the side of the E2s as well.

In order to exclude the possibility that the failure of Pib1 to activate Ubc4 was due to the lack of a suitable substrate, we performed thioester discharge assays with Ubc13 and Ubc4. Here, the ability of an E3 to stimulate E2 activity is measured by its effect on a substrate-independent discharge of the ubiquitin thioester from the E2 to free lysine (Branigan et al, 2015). In order to prevent an intramolecular attack on the catalytic cysteine in Ubc13 or Ubc4, we used E2 mutants, Ubc13(K92R) and Ubc4(K91R), which are catalytically active, but resistant to autoubiquitylation (McKenna et al, 2001). Consistent with its high selectivity for Ubc13, Pib1 promoted the discharge of the Ubc13 thioester (Ubc13~Ub), but not the Ubc4 thioester (Ubc4~Ub), whereas effective discharge of Ubc4~Ub was observed with Rsp5 (Figure 2G, H).

Based on these observations, we conclude that Pib1 acts as an E3 with high selectivity for Ubc13-Mms2, and – as was observed with other RING E3s – the E2 dictates the linkage specificity of the reaction.

### Ubc13-Mms2 cooperates with multiple RING E3s associated with internal membranes

Pib1 shares its domain arrangement with a number of proteins from other organisms (Figure 3A). The FYVE domain in association with a RING finger is conserved among fungi such as *Schizosaccharomyces pombe* and *Ustilago maydis*, whereas FYVE-like domains lacking the characteristic WXXD signature motif can be found in metazoan RING finger proteins such as human Rififylin (RFFL) (Stenmark & Aasland, 1999; Tibbetts et al, 2004). We thus asked whether the selectivity for Ubc13-Mms2-mediated K63-polyubiquitylation would extend to other members of this FYVE-(type)-RING protein family: *S. pombe* SpPib1, *U. maydis* Upa1, and human Rififylin (RFFL). We also analysed two additional RING E3s associated with the endocytic compartment: the integral membrane protein Tul1 from *S. cerevisiae*, which localises to endosomal, Golgi and vacuolar membranes (Li et al, 2015; Reggiori & Pelham, 2002), and human ZNRF2, which lacks a FYVE domain, but is targeted to internal membranes via myristoylation and was reported to be an orthologue of SpPib1 (Araki & Milbrandt, 2003; Wood et al, 2012).

**Figure 3.**
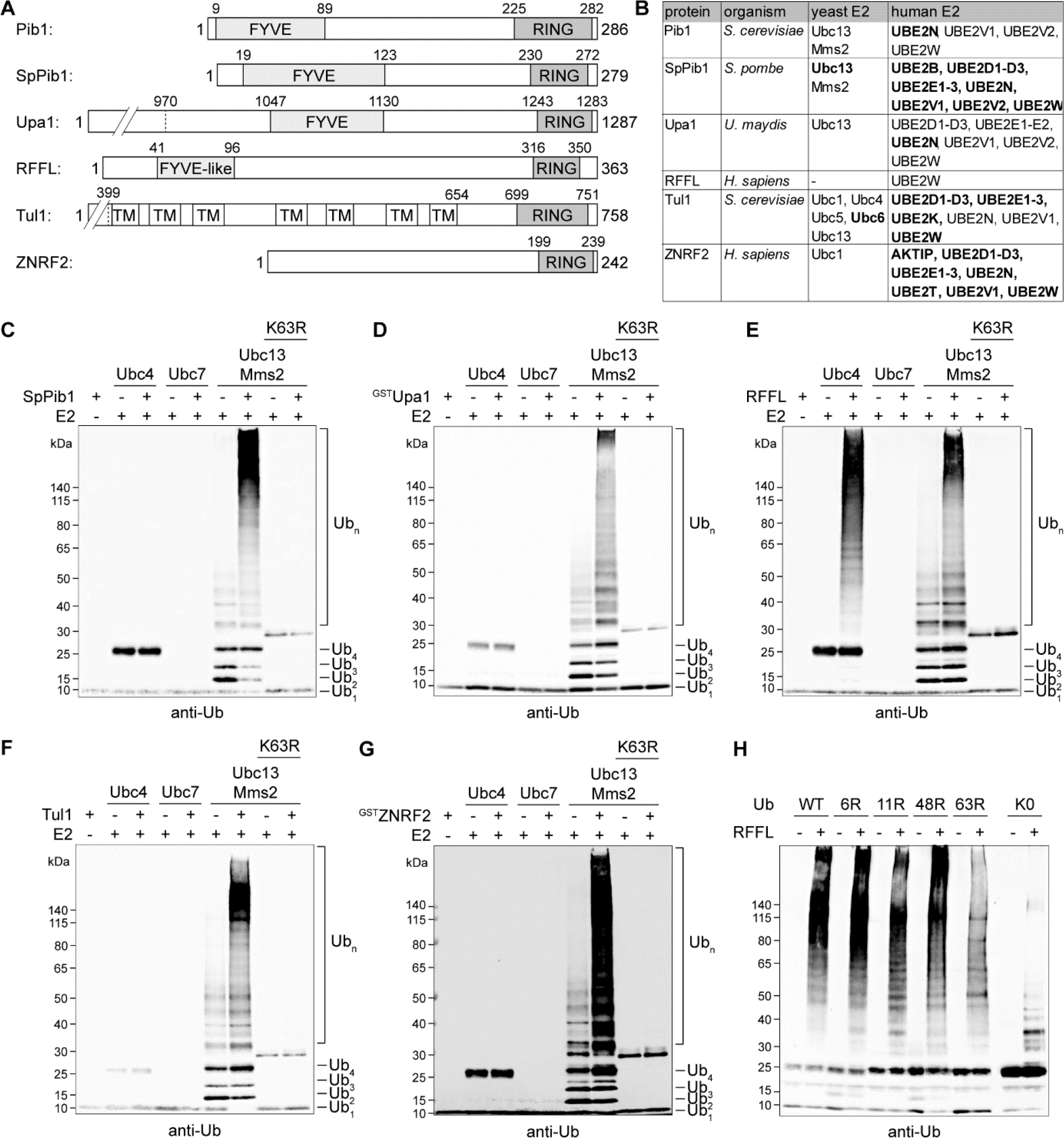
Other membrane-associated RING E3s share Pib1’s selectivity for Ubc13-Mms2. **(A)** Domain structures of the indicated membrane-associated RING E3s (TM: transmembrane domain). **(B)** Membrane-associated RING E3s interact with a range of budding yeast and/or human E2s in the two hybrid assay. Strong interactions are indicated in bold. Original plates are shown in Figure S4. **(C-G)** Selected RING E3s cooperate with Ubc13-Mms2 in K63-polyubiquitylation. Recombinant SpPib1(123-279) **(C)**, ^GST^Upa1(1131-1287) **(D)**, RFFL(97-363) **(E)**, Tul1(655-758) **(F)** and ^GST^ZNRF2 **(G)** were analysed in ubiquitin chain formation assays with the indicated recombinant E2s (see Figure 2D-F) as described in Figure 2A. All reactions contained E1, ATP and ubiquitin (WT or K63R) and were carried out at 30°C for 60 min. **(H)** RFFL-stimulated polyubiquitylation by Ubc4 results in heterogeneous linkages at multiple sites. Polyubiquitylation assays were performed as in (C-G) with WT ubiquitin and indicated lysine mutants.

Systematic two-hybrid assays (Figure 3B, Figure S3A, B) revealed interactions of the fungal FYVE-RING proteins with budding yeast Ubc13 and its human homologue (UBC13 or UBE2N) and weaker interactions with the Ubc13 cofactors, Mms2 and the human E2 variants, UBE2V1 (UEV1A) and UBE2V2 (MMS2). In addition, SpPib1 displayed robust interactions with a range of human E2s, among them members of the UBCH5 family (UBE2D1, UBE2D2 and UBE2D3). A similar, possibly even less selective interaction pattern was observed by ZNRF2. In contrast, RFFL did not show any strong interactions in the two-hybrid assay. The cytoplasmic domain of Tul1 displayed weak interactions with yeast Ubc13 or Mms2, but also with Ubc6 and a number of human E2s, including the UBCH5 family.

Stable associations between E2 and RING finger proteins do not necessarily correlate with efficient cooperation in ubiquitin transfer (Lorick et al, 1999). Therefore, we performed *in vitro* ubiquitin polymerisation assays using the purified E3s in combination with budding yeast Ubc4, Ubc7 and Ubc13-Mms2. All of the RING finger proteins strongly stimulated Ubc13-Mms2 in K63-polyubiquitylation (Figure 3C-G), indicating that robust interactions are not required for productive cooperation. In contrast, none of the E3s stimulated Ubc7 activity. RFFL also promoted the assembly of ubiquitin conjugates by Ubc4. For this E2-E3 pair, use of a series of ubiquitin mutants indicated a diverse set of linkages and attachment sites among the conjugates, as only a lysine-less ubiquitin variant significantly interfered with their assembly (Figure 3H).

In order to exclude the possibility that the observed selectivities were due to species-specific features, we repeated the ubiquitin polymerisation assays with recombinant human E2s, UBC13-UBE2V2 and UBCH5A. As expected, human UBC13-UBE2V2 was active with both human E3s (Figure S3C). Human UBCH5A proved even less selective than its yeast homologue, Ubc4, cooperating at least to some degree with all the constructs tested (Figure S3C). However, stimulation of Pib1 by UBCH5A was marginal compared to Ubc13-Mms2 or UBC13-UBE2V2.

Taken together, these observations suggest that K63-specific polyubiquitylation in cooperation with Ubc13-Mms2 may be a salient feature of this entire group of membrane-associated RING E3s. While some of them also appear to be able to cooperate with other E2s, possibly resulting in alternative linkages, Pib1 is one of the most highly selective E3s.

### Ubc13-Mms2 co-localises with Pib1 at internal membranes

Pib1 localises to membranes of the endocytic compartment via its lipid-binding FYVE domain and accumulates at the vacuolar periphery (Burd & Emr, 1998). As expected, the fungal E3s featuring a genuine FYVE domain localised in a Pib1-like pattern when expressed as fusions to GFP in budding yeast (Figure 4A). In contrast, RFFL was targeted predominantly to the plasma membrane. This is consistent with the situation in mammalian cells, where the protein has been reported to associate with both the plasma membrane and the endocytic compartment (Coumailleau et al, 2004; McDonald & El-Deiry, 2004), and its targeting was described to be in part governed by palmitoylation (Araki et al, 2003).

**Figure 4.**
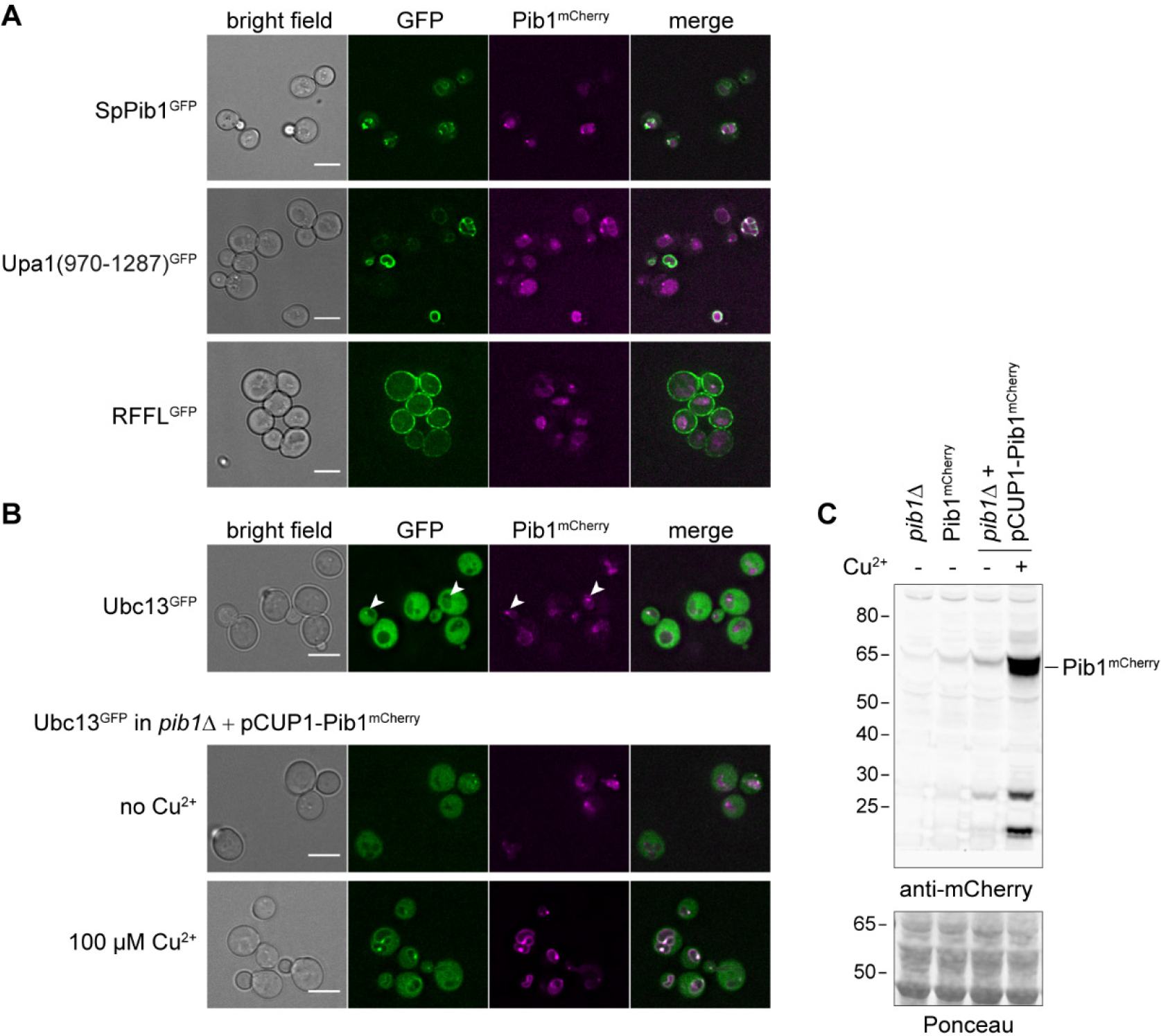
Localisation of RING E3s and Ubc13 in budding yeast cells. **(A)** Selected RING E3s localise in distinct patterns in budding yeast. Cells harbouring endogenously tagged Pib1^mCherry^ and GFP-tagged E3s expressed from a centromeric plasmid under control of the *CUP1* promotor were grown in SC medium containing 70 μM CuSO_4_ at 30°C, and images were acquired by deconvolution microsopy (scale bar = 5 μm). Single representative deconvolved z-planes are shown. **(B)** Pib1 recruits Ubc13 to intracellular membrane structures. Exponentially growing cultures of yeast cells expressing Ubc13^GFP^ and Pib1^mCherry^ in SC medium were imaged by deconvolution microscopy (scale bar = 5 μm). Single representative deconvolved z-planes are shown. Arrow heads highlight co-localisation of Pib1 and Ubc13. Top row: endogenously tagged Pib1^mCherry^; middle row: Pib1^mCherry^ expressed under control of the *CUP1* promoter, in the absence of CuSO_4_; bottom row: Pib1mCherry expressed under control of the *CUP1* promoter, in the presence of 100 μM CuSO_4_. **(C)** Comparison of Pib1^mCherry^ protein levels in the strains used in (B), along with *pib1Δ* harbouring Ubc13^GFP^.

Although the Ubc13-Mms2 complex is mostly distributed in the cytoplasm in a diffuse pattern, sporadic vacuole-associated Ubc13^GFP^ foci co-localising with Pib1^mCherry^ were detectable in some of the cells, consistent with an interaction of the proteins *in vivo* (Figure 4B). Upon overexpression of Pib1^mCherry^ by means of a copper-inducible promoter, localisation of Ubc13^GFP^ to the vacuolar membrane was strongly enhanced, indicating that Pib1 interacts with Ubc13 in cells and is able to recruit the E2 to internal membranes (Figure 4C).

### Genetic interactions implicate Ubc13-Mms2 in membrane protein sorting

The cooperation of Ubc13-Mms2 with the membrane-associated E3s *in vitro* and their co-localisation in cells strongly suggested a functional involvement of the E2 in the endocytic compartment. In order to obtain evidence for this notion, we used genetic interaction profiles as an unbiased approach to identify gene functions (Costanzo et al, 2011). We mined a previously published genome-scale genetic interaction map (Costanzo et al, 2016) for genes whose interaction patterns resembled those of *UBC13* and *MMS2*. This map comprises fitness measurements based on colony size for ~23 million double-mutant combinations, covering roughly 90% of budding yeast genes. In this data set, the genetic interaction profiles of *UBC13* and *MMS2* correlated not only with each other, but also – to a comparable extent – with those of various genes involved in vacuolar transport and the MVB pathway (Figure 5A, B). Based on gene ontology (GO) term analysis, these pathways were strongly overrepresented among the most highly correlated genes for both *UBC13* and *MMS2* (Figure 5C). In particular, numerous genes encoding components of all ESCRT complexes (ESCRT-0: *HSE1*; ESCRT-I: *SRN2*, *MVB12*; ESCRT-II: *VPS25*; ESCRT-III-associated: *VPS4*) emerged at the top of the list. Analysis of individual genetic relationships revealed significant negative interactions of both *ubc13Δ* and *mms2Δ* mutants with candidates that were expected based on their contribution to genome maintenance (such as *rev1Δ*, *rev3Δ*, *rev7Δ* and *pol32Δ*), but also with three different *rsp5* alleles and (for *ubc13Δ*) with *tul1Δ* (Figure 5D, E). Thus, both genetic interaction profiles and individual genetic interactions suggested a function of Ubc13-Mms2 in membrane protein sorting. We therefore asked whether Ubc13 might cooperate with Pib1 in this context.

**Figure 5.**
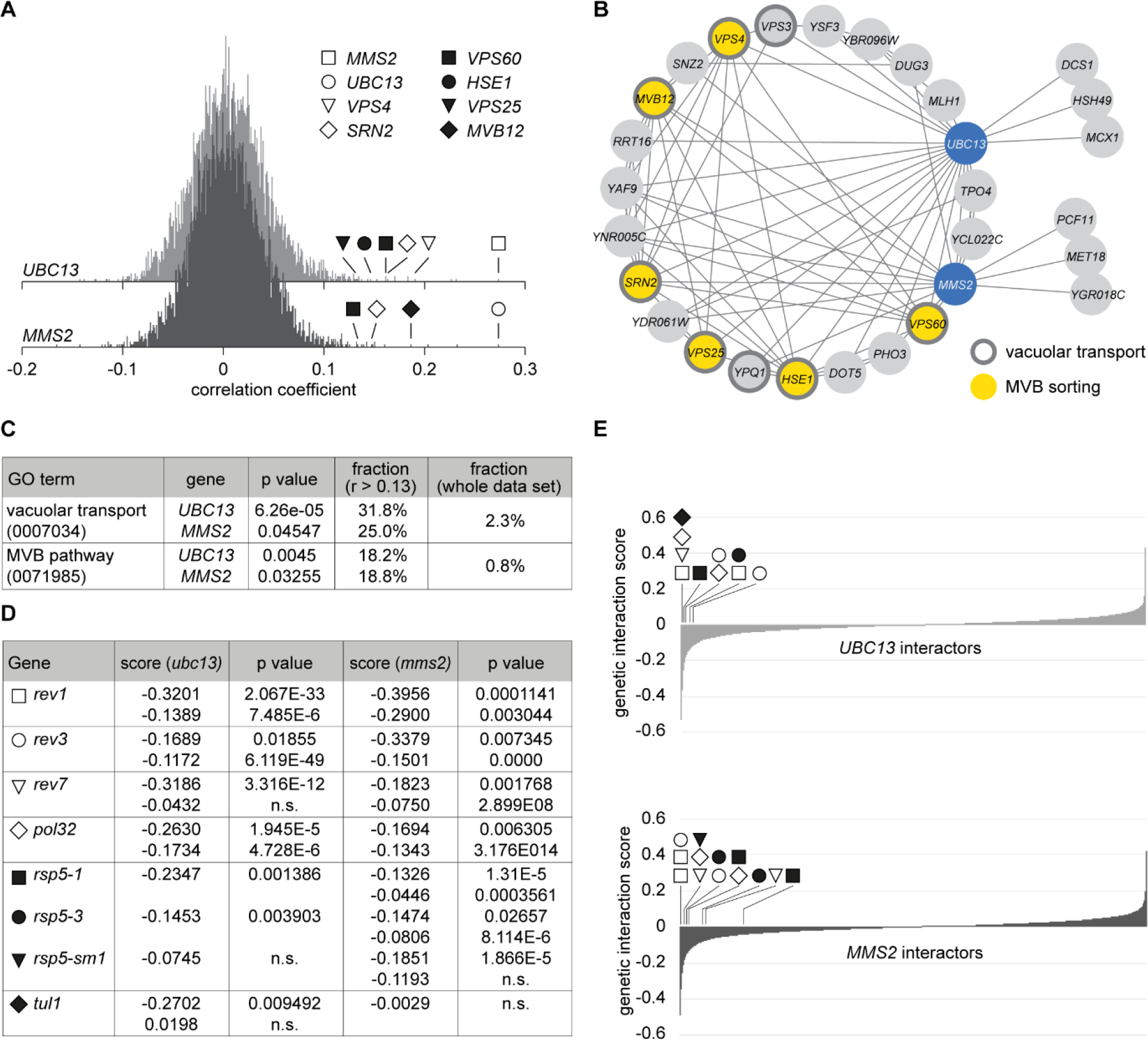
Genetic interactions implicate *UBC13* and *MMS2* in membrane protein sorting. **(A)** Functional similarity between *UBC13*, *MMS2* and components of membrane protein sorting pathways. Histograms of Pearson correlation coefficients calculated between the genetic interaction profiles of *UBC13* and *MMS2* and ~ 90% of all yeast genes are shown, obtained from a previously published genome-scale genetic interaction map (Costanzo et al, 2016). A table of all interactions included in the analysis is provided as Supplementary Material. **(B-C)** Genes involved in vacuolar transport and the MVB pathway are overrepresented among those whose interaction profiles most strongly correlate with *UBC13* and *MMS2* (Pearson correlation coefficient > 0.13). **(B)** Network of genes correlating with *UBC13* or *MMS2*. All correlations within the group above the 0.13 threshold are shown. Genes involved in vacuolar transport (gene ontology term GO: 0007034) and the MVB sorting pathway (GO: 0071985) are highlighted. **(C)** Fraction of genes with relevant GO terms among those depicted in (B) versus the total fraction of genes with these GO terms in the data set. Significance is indicated by p-values. **(D-E)** *UBC13* and *MMS2* exhibit negative genetic interactions with genes involved in translesion synthesis and membrane protein sorting. **(D)** Genetic interaction scores of *UBC13* and *MMS2* with relevant genes, obtained from the same genetic interaction map as in (A). Significance is indicated by p-values (n.s.: not significant, p>0.05). Note that two interaction scores are provided for some combinations, based on the availability of query and array alleles in the library. **(E)** Plots of all genetic interaction scores for *UBC13* and *MMS2*. Symbols indicate significant interactors from (D).

### Ubc13-Mms2 contributes to endocytosis and the MVB pathway

The v-SNARE Snc1 was previously found to be subject to K63-linked polyubiquitylation (Xu et al, 2017). This modification was reported to mediate interactions with the COPI complex, thus contributing to the recycling of Snc1 to its donor membranes (Xu et al, 2017). Based on the impairment of a *pib1Δ tul1Δ* double mutant in Snc1 recycling, Pib1 and Tul1 were postulated to act redundantly as E3s on the v-SNARE (Xu et al, 2017). In this case, Ubc13-Mms2 could thus contribute to the modification as an E2. However, although Snc1 ubiquitylation was detectable in our hands, the modification pattern was not altered in *pib1Δ*, *tul1Δ* or *pib1Δ tul1Δ* mutants (Figure S4A). Snc1 had also been identified as a target of K63-ubiquitylation upon treatment of yeast with hydrogen peroxide in a proteomic screen (Silva & Vogel, 2015). Although Snc1 ubiquitylation levels were enhanced by H_2_O_2_, deletion of *PIB1* and/or *TUL1* did not diminish them (Figure S4B). These results argue against Pib1 and Tul1 acting as cognate E3s for Snc1, but rather imply an indirect effect on its recycling or turnover.

In the MVB pathway, Pib1 acts redundantly with an endosomal Rsp5 adaptor, Bsd2, which facilitates transport of endocytic vesicles from the Golgi to the vacuole (Nikko & Pelham, 2009). In consequence, plasma membrane proteins such as the uracil permease (Fur4) accumulate in endosomes of *bsd2Δ pib1Δ* double mutants and fail to be degraded in the vacuole. Thus, if Ubc13-Mms2 cooperate with Pib1, then *ubc13* and *pib1* mutations should behave epistatically with respect to Fur4 degradation in a *bsd2Δ* background. We therefore measured the vacuolar degradation of GFP-tagged Fur4 induced in an established procedure by cycloheximide treatment (Volland et al, 1994). As GFP is not degraded but accumulates in the vacuole, we followed the appearance of free GFP as a reporter for vacuolar delivery (Figure 6A). This approach allows quantification of an otherwise entirely microscopy-based assay (Nikko & Pelham, 2009) (Figure 6B). A *doa4Δ* mutant served as a control as it was previously shown to be completely defective in Fur4 internalisation and thus vacuolar targeting (Galan & Haguenauer-Tsapis, 1997). As reported previously (Nikko & Pelham, 2009), deletion of *PIB1* had no effect in isolation, but impeded Fur4^GFP^ degradation in a *bsd2Δ* background. The *tul1Δ* mutant or combinations of *tul1Δ* with *bsd2Δ* or *pib1Δ* were not impaired in Fur4 degradation, suggesting that Tul1 does not functionally overlap with Pib1 in Fur4 degradation (Figure S4C). Notably, however, the *ubc13Δ* mutant alone exhibited a phenotype similar and additive to that of *bsd*2*Δ* (Figure 6A, B). Importantly, *pib1Δ* did not significantly enhance the *ubc13Δ* phenotype, either alone or in combination with *bsd2Δ.* This implies that Ubc13-Mms2 cooperates with Pib1 in vacuolar delivery of Fur4.

**Figure 6.**
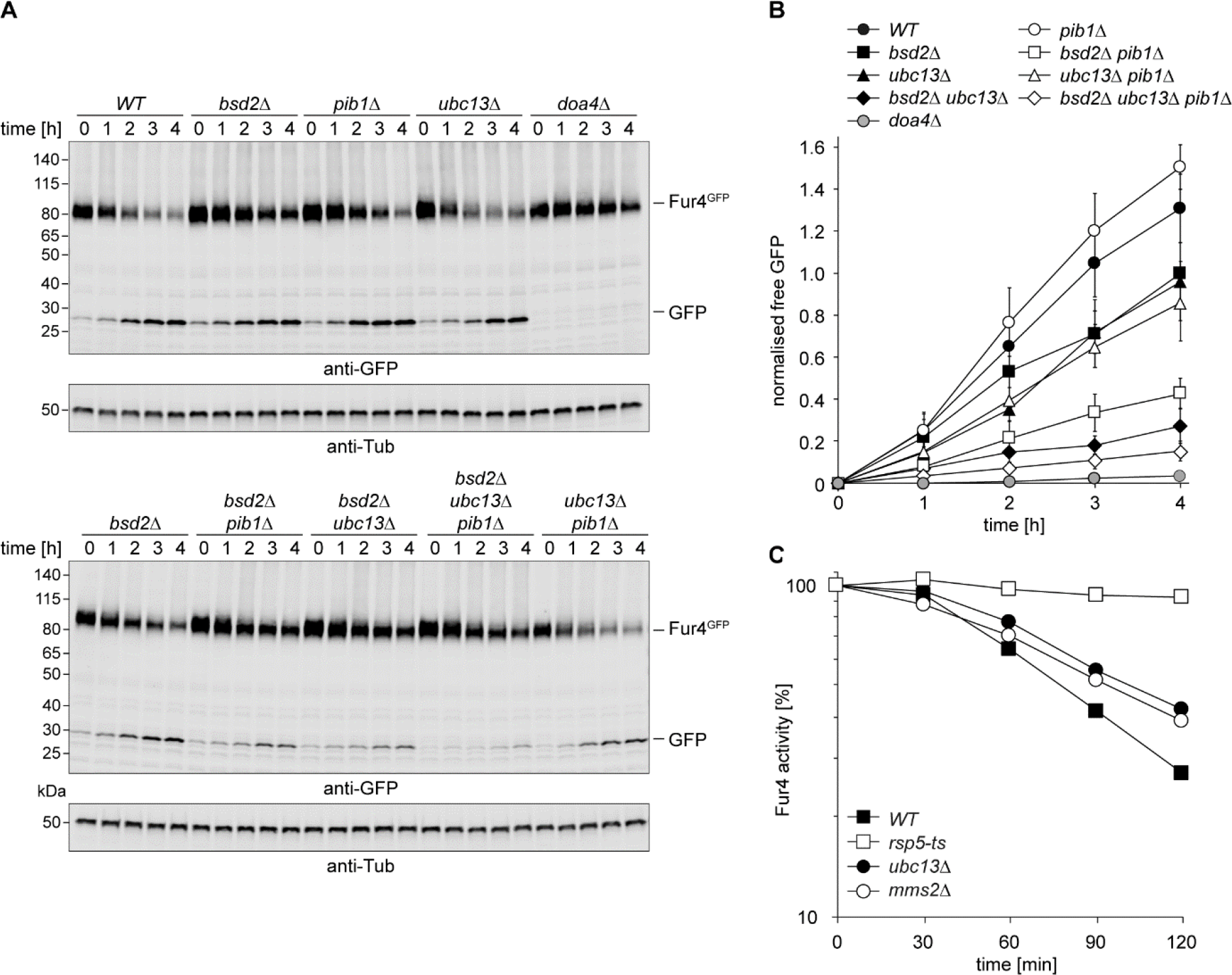
Ubc13-Mms2 contributes to membrane protein sorting. **(A)** Pib1 and Ubc13 cooperate in the delivery of Fur4 to the vacuole. Internalisation of Fur4^GFP^, expressed from a centromeric plasmid under control of a *GAL1* promotor, was induced in strains of the indicated backgrounds by addition of 100 μg/ml cycloheximide. Samples were taken at the indicated times, and total cell extracts were analysed by western blotting with anti-GFP and anti-α-tubulin antibodies. **(B)** Quantification of vacuolar degradation of Fur4^GFP^. Accumulation of free GFP on the blots shown in (A) was plotted relative to the GFP signal at 4 h in *bsd2Δ*. Error bars indicate standard deviations derived from three independent experiments with two technical replicates each. The α-tubulin signal was used as a loading control. For details, see Materials and Methods. **(C)** Internalisation of Fur4 is delayed in *ubc13Δ* and *mms2Δ*. Fur4^myc^ was overexpressed from a multicopy plasmid, and internalisation of the permease was induced by cycloheximide treatment as in (A-B). The amount remaining at the plasma membrane at the indicated time points was quantified relative to t=0 by uracil uptake assays. A temperature-sensitive *rsp5* mutant served for comparison.

The defect of the *ubc13Δ* single mutant suggested that the E2 fulfils additional roles in membrane trafficking not shared by Pib1 or Bsd2. Indeed, when we monitored the internalisation of Fur4 from the plasma membrane by uracil uptake assays after cycloheximide treatment (Volland et al, 1994), we observed delays in both *ubc13Δ* and *mms2Δ* (Figure 6C). These were not due to impaired ubiquitylation of the permease itself (Figure S4D), consistent with previous reports ascribing Fur4 modification to Rsp5 (Galan et al, 1996). Ubc13-mediated ubiquitylation therefore appears to exert an effect on the endocytic system that is distinct from Rsp5, but important for the uptake of plasma membrane proteins like Fur4.

In conclusion, our observations suggest that K63-polyubiquitylation in the endocytic compartment is mediated not exclusively by Rsp5, but also by Ubc13-Mms2. Like Rsp5, the E2 appears to contribute to more than one aspect of membrane trafficking, affecting both internalisation of a plasma membrane protein and – in cooperation with Pib1 – delivery of vesicles to the vacuole via the MVB pathway.

## Discussion

### Contribution of Ubc13-Mms2 to K63-polyubiquitylation at internal membranes

Polyubiquitin chains of K63-linkage have been implicated in many aspects of membrane trafficking (Erpapazoglou et al, 2014). In budding yeast, Rsp5 has so far been the only verified source of that chain type within the cytoplasmic compartment in yeast. Our results now show that Ubc13-Mms2, aided by the RING E3 Pib1 and potentially other membrane-associated E3s, contributes to protein sorting in the MVB pathway, thus expanding the scope of this E2 beyond its known nuclear function with Rad5 in genome maintenance.

Recombinant Pib1 was previously reported to cooperate *in vitro* with a highly non-selective E2, UBCH5A (Shin et al, 2001), which is not known for a particular bias towards the K63-linkage. Our results now indicate a clear preference of Pib1 for Ubc13-Mms2, thus strongly implicating the E2-E3 pair in the formation of K63-chains at internal membranes. An earlier claim about K63-specificity of Pib1 was made by Xu *et al*. (Xu et al, 2017), who observed a redundant function of Pib1 and Tul1 in COPI-mediated vesicle sorting. They found that an interaction of the COPI complex with a ubiquitylated v-SNARE, Snc1, mediated by K63-selective ubiquitin binding, drives the recycling of the v-SNARE to its donor membranes. Deletion of *PIB1* and *TUL1* inhibited Snc1 recycling, and fusion of a non-specific deubiquitylating enzyme (DUB) to Pib1 or Tul1 interfered with Snc1 ubiquitylation and trafficking (Xu et al, 2017). From these indirect data they concluded that Pib1 – together with Tul1 – acts as a cognate E3 in K63-ubiquitylation of Snc1. However, by direct observation of its modification pattern we found no defect in Snc1 ubiquitylation in the *pib1Δ tul1Δ* mutant, which suggests an indirect effect of the ligases on vesicle trafficking rather than an involvement in ubiquitin conjugation to the v-SNARE. The observed dominant negative effects exerted by the E3-DUB fusions could be attributable to a physical proximity of the ligases to Snc1 mediated by the E3s’ membrane association, but they do not prove an enzyme-substrate relationship.

Our findings now provide conclusive evidence for a cooperation of Ubc13-Mms2 with Pib1 and possibly additional E3s in K63-polyubiquitylation at endosomal and vacuolar membranes. However, they do not reveal any direct physiological substrates of these enzymes. It is well established that cellular membrane trafficking, particularly in the endocytic pathway, requires multiple ubiquitylation events at various stages and on many different targets, and emerging information about E3s like Rsp5, Pib1 and Tul1 suggests a high degree of redundancy in the system. Moreover, the exquisite specificity of Ubc13-Mms2 for K63 of ubiquitin could indicate a predominant function in chain elongation rather than *de novo* modification of specific substrates, as observed in cooperation with Rad5 (Hoege et al, 2002; Ulrich & Jentsch, 2000).

### Selectivity of E2-E3 pairing

Our identification of budding yeast Pib1 as an E3 with an unusually high preference for the K63-specific Ubc13-Mms2 complex raises the general question about the origin of selectivity in E2-E3 cooperation. Based on sequence or structural information alone, predictions about productive pairings have not yet been successful, necessitating a combination of biochemical and functional experiments to address this issue. The reported co-localisation and redundancy between Pib1 and Tul1, along with our *in vitro* activity assays, suggest that Tul1 – like Pib1 – cooperates with Ubc13-Mms2 in K63-ubiquitylation. Yet, the protein was originally described to interact with Ubc4 (Reggiori & Pelham, 2002). Although we were unable to reproduce this finding or detect any catalytic activity with yeast Ubc4, our interaction and activity assays suggest that Tul1 might be less selective than Pib1.

Ubc13-Mms2 was also found to be active with a family of related E3s from other species, all harbouring a lipid-binding FYVE- or FYVE-like domain in addition to the RING finger. Interestingly, while they all supported Ubc13-Mms2 activity, the scope of their interactions and activities with other E2s varies considerably. The putative Pib1 homologue from *S. pombe*, SpPib1, displayed an identical localisation and a very similar activity pattern, although its interactions in the two-hybrid system were less selective than those of budding yeast Pib1. Given its preference for Ubc13, it is surprising that SpPib1 was reported to target a subunit of the exocyst, Sec3, for proteasomal degradation (Kampmeyer et al, 2017), a process usually associated with K48-linked polyubiquitylation. However, a degree of redundancy between Pib1 and other E3s (including Rsp5) was noted in that study, and a contribution of vacuolar degradation was not rigorously excluded, thus leaving open the possibility that multiple degradation systems act on Sec3. *U. maydis* Upa1 closely resembled budding yeast Pib1 with respect to localisation, E2 binding and E2 activation, implying a similarly Ubc13-selective function. Interestingly, the protein bears a large N-terminal extension, including an RNA-binding domain that implicates Upa1 in vesicle-mediated long-distance transport of mRNP particles (Pohlmann et al, 2015). Thus, the protein appears to couple endosome dynamics with RNA transport.

RFFL (also known as CARP2 or Sakura) displayed the lowest selectivity among the FYVE-type E3s. Accordingly, the protein has been reported to modify a diverse set of substrates, many of them localising to the plasma membrane or endocytic vesicles (Araki et al, 2003; Liao et al, 2008; McDonald & El-Deiry, 2004; Okiyoneda et al, 2018). RFFL was often observed to act redundantly with a closely related protein, RNF34 (also called CARP1 or Momo) (Araki et al, 2003; Liao et al, 2009). Whereas most instances of RFFL- and/or RNF34-mediated ubiquitylation appear to involve proteasomal degradation (Araki et al, 2003; Liao et al, 2009; Liao et al, 2008; McDonald & El-Deiry, 2004; Wei et al, 2012; Yang et al, 2007; Zhang et al, 2014), there are cases where a contribution of lysosomal proteolysis has been demonstrated (Jin et al, 2014; Okiyoneda et al, 2018). For example, in a recent report RFFL was shown to mediate retrieval from the plasma membrane and subsequent lysosomal degradation of a non-native CFTR mutant via interaction and K63-polyubiquitylation in a post-Golgi compartment. (Okiyoneda et al, 2018). In this context, cooperation with UBC13 appears plausible.

In conclusion, our data point to the existence of a conserved group of membrane-associated RING E3s that can cooperate with Ubc13-Mms2 in the generation of K63-linked polyubiquitin chains and contribute to protein sorting in the endocytic and MVB pathways. The molecular features determining their selectivity as well as the relevant targets of ubiquitylation remain to be explored.

## Materials and Methods

### Yeast strains and plasmids

Yeast strains used in this study are listed in Table S1. Single gene deletions in the BY4741 background were obtained from the yeast knockout collection (Dharmacon). Other gene deletions were generated by PCR-based methods (Janke et al, 2004) or genetic crosses. Tagging of Pib1 and Ubc13 at their endogenous loci was achieved by recombination-mediated integration of PCR-amplified cassettes (Janke et al, 2004). Yeast strains were grown in YPD or, if selecting for maintenance of centromeric or episomal plasmids, synthetic complete (SC) medium lacking the appropriate amino acids.

All plasmids are listed in Table S2. For Fur4 internalisation assays, Fur4^yeGFP^ was expressed under control of a *GAL1* promotor from a centromeric vector based on YCplac33 (Gietz & Sugino, 1988). Two-hybrid plasmids were based on pGBT9 (Clontech) or the pGAD-C- and pGBD-C-series, expressing candidates as fusions to the Gal4 activation domain (GAD) or DNA-binding domain (GBD) and carrying the *LEU2* and *TRP1* markers, respectively (James et al, 1996). For localisation studies of Pib1 orthologues, the corresponding genes were cloned in frame with yeGFP under control of a *CUP1* promotor into a YCplac33-based centromeric plasmid. Overexpression of mCherry-tagged *PIB1* was achieved in the same manner. Plasmids for expression of ^*3HA*^*SNC1* and ^*3HA*^*SNC1(8KR)* under control of the *TPI1* promotor in budding yeast were kindly provided by Todd Graham (Xu et al, 2017). Point mutations were created by PCR-based mutagenesis.

### Yeast two-hybrid assays

The initial two-hybrid screen for interactors of Mms2 was performed with a histidine- and adenine-responsive reporter strain, PJ69-4a, and a set of three yeast genomic libraries kindly provided by Philipp James (James et al, 1996). These libraries, constructed in the pGAD-C vector series representing all 3 reading frames, were amplified in *E. coli*, yielding between 80 and 200 million clones, and used in parallel for transformation of PJ69-4a expressing GBD-Mms2 from a bait plasmid carrying the *URA3* marker. Approximately 5 million clones per library were screened on SC Ura^−^Leu^−^His^−^ selective medium. False positives were eliminated by excluding clones that allowed reporter-dependent growth without the bait plasmid, and genuine interactors were subsequently identified by plasmid isolation and sequencing after re-streaking single colonies.

Individual two-hybrid assays were performed in the haploid strain PJ69-4a as described previously (James et al, 1996). The screen for identifying E2-E3 interaction pairs was performed in a semi-automated manner in a 16×24 spot array format, using a ROTOR HDA pinning robot (Singer Instruments) with a collection of 30 human E2s, a kind gift from Rachel Klevit (Christensen & Klevit, 2009) and all yeast E2s in pGAD-vectors. Open reading frames of budding yeast Pib1, human RFFL (isoform 1, from HeLa cDNA), *S. pombe* Pib1 (aa 123-279), *U. maydis* Upa1 (aa 970-1287) and Tul1 (aa 655-758) were PCR-amplified and cloned into pGBD-C1. Human ZNRF2 was obtained from the Orfeome clone collection (ID 47920) and transferred by Gateway cloning into pGBT9-GW (Clontech). GAD-E2 and GBD-E3 constructs were combined by transformation into PJ69-4α and PJ69-4a, respectively, and subsequent mating. The diploids were then subjected to standard two-hybrid analysis by pinning onto SC Leu^−^Trp^−^, SC Leu^−^Trp^−^His^−^ and SC Leu^−^Trp^−^His^−^Ade^−^ plates. To reduce background growth, the SC Leu^−^ Trp^−^His^−^ plate was re-pinned once onto the same medium. For analysing interactions between Pib1 and the yeast E2s, this screen was repeated in the opposite direction, swapping GAD and GBD.

### Protein purification

Recombinant proteins were produced in *E. coli*. Expression and purification conditions, including buffers, for all recombinant proteins are listed in Table S3. Briefly, His_6_- and GST-tagged proteins were affinity-purified in batch on Ni-NTA agarose (Qiagen) and glutathione Sepharose 4 Fast Flow (GE Healthcare), respectively. Protease cleavage (PreScission: GE Healthcare; TEV-Protease: Biomol) was performed overnight at 4°C. Subsequent purification steps were carried out on an NGC chromatography system (Bio-Rad). Bovine ubiquitin (Ub) was purchased from Sigma Aldrich and Ub(K6R) from BostonBiochem. ^His^Uba1 (from mouse) and other ubiquitin variants were purified as described previously (Carvalho et al, 2012; Pickart & Raasi, 2005). ^GST^Cue1^His^ was purified as described before (Bagola et al, 2013) with some modifications.

### In vitro protein-protein interaction assays

*In vitro* interaction studies were carried out in a buffer containing 50 mM potassium phosphate, pH 6.0, 150 mM KCl, 10% glycerol, 0.1% Triton X100 and 1 mM dithiothreitol (DTT). Purified GST and ^GST^Pib1ΔFYVE (100 pmol) were immobilised on glutathione Sepharose 4 Fast Flow (25 μl slurry, GE Healthcare). Beads were pre-treated with 1 mg/ml bovine serum albumin (Sigma), washed and incubated with 100 pmol His_6_-tagged E2s at 4°C for 120 min. After washing the beads, bound proteins were eluted by boiling in NuPage LDS sample buffer (Thermo Fisher) supplemented with 25 mM DTT for 10 min at 95°C, and the eluate was analysed by SDS-PAGE and western blotting with anti-His_6_-tag antibody (Sigma).

### Ubiquitin chain formation assays

Free ubiquitin chain formation was assayed in reactions containing 40 mM HEPES, pH 7.4, 8 mM magnesium acetate, 50 mM NaCl, 5 μM purified bovine ubiquitin (Sigma) or ubiquitin variants, 50 nM ^His^Uba1, 30 μM ATP, 2 μM E2 and 0.5 μM E3. Reactions were incubated for 30 min (Pib1) or 60 min (other E3s) at 30 or 37°C, stopped by addition of NuPage LDS sample buffer supplemented with DTT, and analysed by SDS-PAGE and western blotting with a monoclonal anti-ubiquitin antibody, P4D1 (Cell Signalling Technology). For subsequent treatment with AMSH (McCullough et al, 2004), the ubiquitylation reaction was terminated by addition of 20 mM EDTA and further incubated in the presence or absence of 2 μM ^GST^AMSH at 37°C for 60 min.

### E2 thioester discharge assays

Ubc13(K92R) was charged with ubiquitin for 60 min at 30°C in a reaction containing 40 mM HEPES, pH 7.4, 2 mM MgCl_2_ 50 mM NaCl, 240 nM His_6_-E1, 30 μM Ubc13(K92R), 24 μM ubiquitin and 200 μM ATP. The reaction was stopped by addition of 1 unit apyrase (New England Biolabs) per 100 μl reaction volume and incubation at 30°C for 10 min (sample t=0). The reaction, now containing the Ubc13~Ub thioester, was mixed with a solution containing L-lysine (adjusted to pH 7.0, Sigma) as well as Mms2 and Pib1−RING+100aa where indicated. Final concentrations were 15 μM Ubc13/Ubc13~Ub, 15 μM Mms2, 1.5 μM Pib1−RING+100aa, 12 μM ubiquitin and 45 mM L-lysine. Reactions were incubated at 30°C for the indicated times, stopped by addition of NuPage LDS sample buffer without DTT and separated by SDS-PAGE, followed by staining with Instant Blue (Expedeon). Ubc4(K91R) was charged with ubiquitin under identical conditions at 30°C for 10 min, and lysine discharge assays were performed with 1.5 μM Pib1−RING+100aa or Rsp5 at a final concentration of 5 mM L-lysine.

### Fluorescence microscopy

Fluorescence microscopy images were acquired with a DeltaVision Elite system (softWoRX 6.5.2) equipped with a 60x objective (NA 1.42) and a DV Elite sCMOS camera (GE Healthcare). Cells were grown to exponential phase and visualised without fixation. Images were deconvolved with softWoRx version 6.5.2 (GE Healthcare) and processed with FIJI (Schindelin et al, 2012).

### Fur4 internalisation and vacuolar delivery assays

Delivery of Fur4 to the vacuole was assayed as described (Nikko & Pelham, 2009). Briefly, yeast cells harbouring YCplac33-GAL-Fur4-GFP were grown to exponential phase (OD_600_ 0.3-0.4) in SC Ura-containing 2% raffinose/0.2% glucose, then shifted to medium containing 2% galactose/0.1% glucose. After 2.5 h, 2% glucose was added to stop Fur4-yeGFP expression, and endocytosis was induced by addition of 100 μg/ml cycloheximide. Samples were collected at the indicated time points and flash-frozen on dry ice. Total cell extracts were prepared by trichloroacetic acid (TCA) precipitation as described previously (Morawska & Ulrich, 2013) with some modifications: 1. precipitation was performed directly in SC medium without pelleting the cells; 2. the final pellet was resuspended in NuPage LDS sample buffer supplemented with 25 mM DTT and incubated for 20 min at 30°C. Samples were analysed by SDS-PAGE followed by western blotting with mouse anti-GFP (Roche) and rabbit anti-α-tubulin antibodies (Abcam).

Western blots were imaged with an Odyssey CLx system (LI-COR) using near-infrared fluorophore-labelled secondary antibodies. Signals were quantified (background correction: average) with Image Studio Version 3.1 (LI-COR). For quantification of vacuolar degradation of Fur4, three independent biological replicates with two technical replicates each were performed. For each sample, the intensity of free GFP (800 nm channel) was normalised by division by the intensity of the tubulin loading control (700 nm channel). To account for variations in the amount of free GFP at t=0 in a given strain, these values were corrected by subtracting the normalised intensity at t=0. The resulting corrected signal of GFP at 4 h in the *bsd2Δ* strain was arbitrarily set to 1 for each individual western blot, and the other values were calculated relative to this reference point.

Fur4 internalisation was measured by uracil uptake assays as described (Galan & Haguenauer-Tsapis, 1997; Volland et al, 1994). Briefly, exponential cultures of the relevant strains expressing Fur4^myc^ from a multicopy plasmid were treated with 100 μg/ml cycloheximide. At the indicated time points, 1 ml aliquots were incubated with 5 μM [^14^C]uracil for 20 s at 30°C and filtered immediately through Whatman GF/C filters. Filters were then washed twice with ice-cold water and subjected to scintillation counting. The amount of radiation taken up at each time point was plotted relative to the value at t=0.

### Detection of Fur4 and Snc1 ubiquitylation

Fur4 ubiquitylation was detected by expression of Fur4^myc^ from a multicopy plasmid in the relevant yeast strains, preparation of a membrane-enriched lysate from exponential cultures as described (Galan et al, 1996), SDS-PAGE and western blotting with an anti-myc antibody (9E10). For detection of Snc1 ubiquitylation, relevant yeast strains expressing wild-type 3HA-tagged Snc1 (pRS416-3xHA-Snc1) or a lysineless mutant (pRS416-3xHA-Snc1(8KR)) were grown to exponential phase, treated – where indicated – with 2.4 mM H_2_O_2_ for 45 min at 30°C and subjected to total cell extraction with TCA (see above), followed by SDS-PAGE and western blot analysis using an anti-HA-antibody (Santa Cruz).

## Supporting information

UBC13 MMS2 genetic interaction data

## Acknowledgements

We thank S. Braun, K. Schmidt and M. Berkenkopf for assistance with experiments, H.-P. Wollscheid and S. Wegmann for sharing biochemical expertise, and IMB’s media laboratory for supplies. We are grateful to J. Azevedo, M. Ellison, M. Feldbrügge, J. Huibregtse, P. James, T. Graham, R. Klevit, D. Komander and T. Sommer for supplying reagents and advice. This work was funded by grants from the European Research Council to H.D.U. (ERC AdG 323179 and ERC PoC 786330).

## Author Contributions

C.R. and H.D.U. conceived the study, C.R., V.T., T.K.A., O.S., S.J. and H.D.U. performed and analysed experiments; A.K. analyzed genetic interaction profiles; H.D.U. wrote the manuscript with input from C.R.; C.R., A.K. and H.D.U. prepared the figures, and V.T., T.K.A., O.S. and A.K. commented on the manuscript.

## Supplementary Material

**Figure S1.**
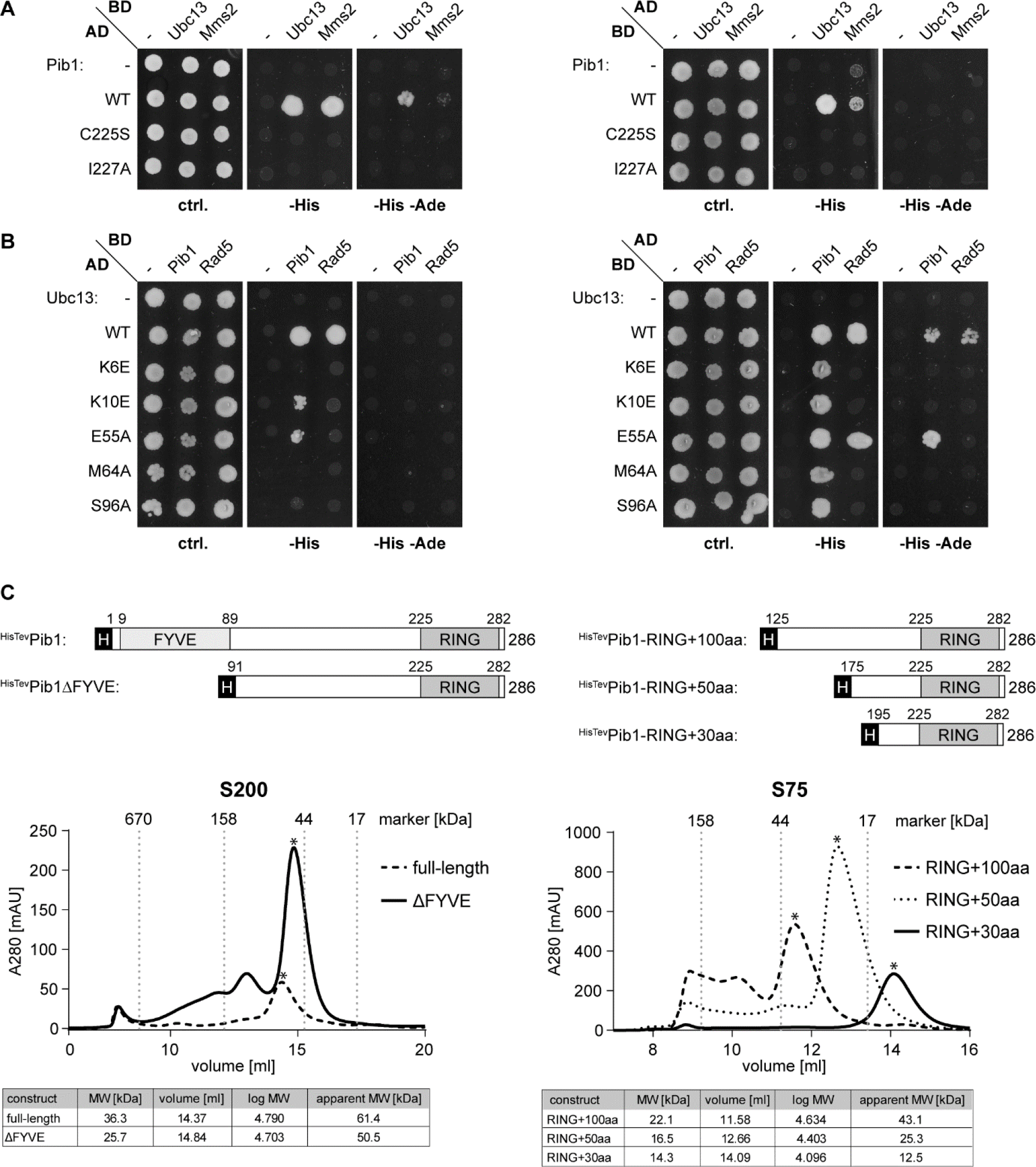
Protein-protein interactions of Pib1, Ubc13 and Mms2. **(A)** Mutations in the Pib1 RING domain abolish interactions with Ubc13 and Mms2 in the two-hybrid system. Assays were performed as described in Figure 1B. **(B)** Mutations in Ubc13 that abolish interaction with Rad5 also affect interaction with Pib1. Two-hybrid assays were performed as described in Figure 1B. **(C)** Pib1 dimerisation requires a domain N-terminal of its RING finger. Domain structures (numbering according to Pib1; H: His_6_-Tev tag) and gel filtration profiles of the indicated recombinant His_6_-TEV-tagged Pib1 constructs, expressed in *E. coli* and purified by Ni-NTA affinity chromatography. Profiles were recorded on calibrated (Gel Filtration Standard, BioRad) Superdex 200 or Superdex 75 10/300 GL columns. Asterisks indicate the relevant peaks. Tables beneath the elution profiles indicate elution volume and apparent as well as actual molecular weights.

**Figure S2.**
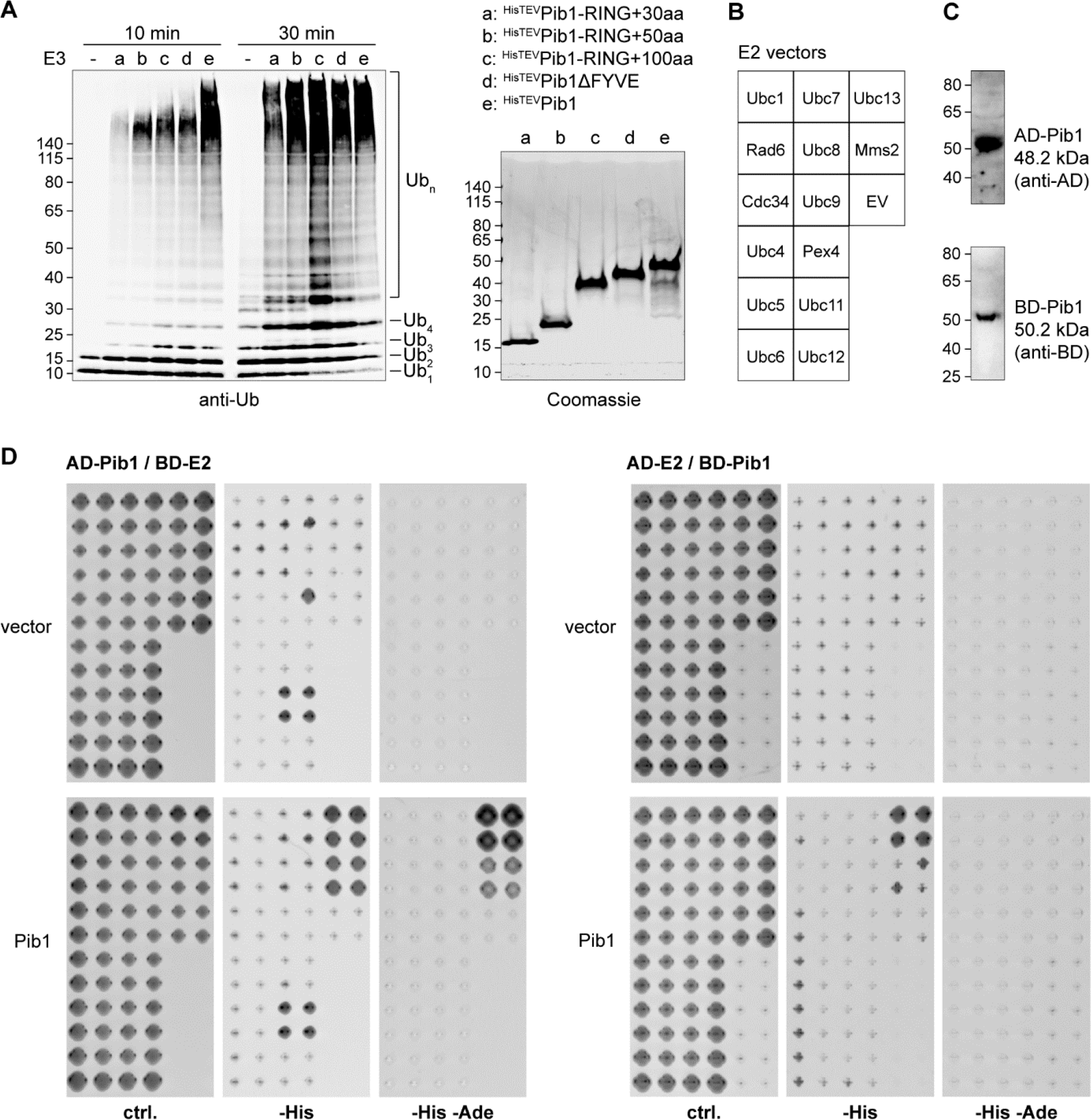
Pib1 is a cognate E3 for Ubc13-Mms2. **(A)** Pib1 is most active as a dimer. *In vitro* polyubiquitin chain formation assays were carried out as described in Figure 2A, using the indicated His_6_-TEV-tagged constructs. Reaction products are shown on the left (anti-ubiquitin blot); the right-hand panel shows a Coomassie-stained gel of the relevant E3 constructs. **(B-D)** Among all budding yeast E2s, only Ubc13-Mms2 interacts with Pib1 in the two-hybrid system. Systematic two-hybrid interaction assays of Pib1 with all budding yeast E2s (and Mms2) were performed in diploid cells, generated by mating haploid a and α cells expressing the indicated Gal4 activation domain (AD) fusion constructs and Gal4 DNA-binding domain (BD) fusion constructs, respectively. Selected diploid cells were spotted as indicated in the scheme (EV: empty vector) **(B)**. Expression of Pib1 constructs was verified by western blotting of total cell extracts with anti-AD and anti-BD antibodies **(C)**, and cultures were subjected to growth analysis on selective plates **(D)**.

**Figure S3.**
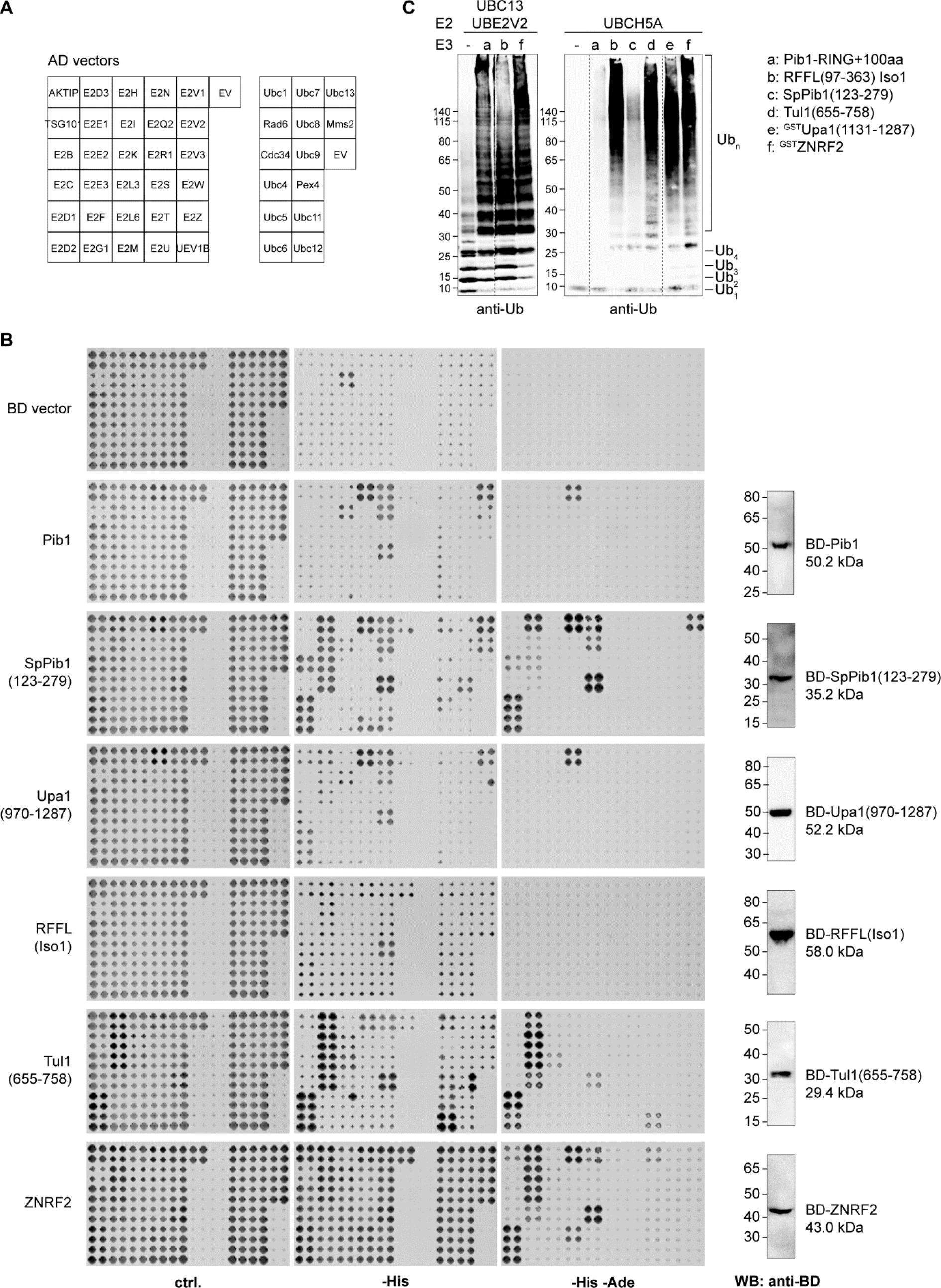
Interactions and activities of selected membrane-associated E3s with E2s. **(A)** Spotting pattern of the human and budding yeast AD-E2 constructs for systematic two-hybrid assays (EV: empty vector). **(B)** Left: Systematic two-hybrid assays of relevant E3s with 30 human E2s and E2-like proteins and all budding yeast E2s, performed as described in Figure S2B-D. Right: images of the corresponding BD-E3 constructs, derived from western blots of total cell extracts using an anti-BD antibody. Note that the panels showing interactions of BD-Pib1 with yeast AD-E2s and the corresponding expression control are identical to those shown in Figures S2C and D, and are included here again for comparison. **(C)** Human E2s can substitute for yeast Ubc13-Mms2 and Ubc4. Left: Pib1 and human E3s stimulate polyubiquitin chain synthesis by human UBC13-UBE2V2. *In vitro* polyubiquitylation assays were performed as described in Figure 3C-G with the indicated E3 constructs, using UBC13-^His^UBE2V2 as E2. Right: UBCH5A activates E3s in a highly non-selective manner. *In vitro* polyubiquitylation assays were performed as described in Figure 3C-G with the indicated E3 constructs, using ^His^UBCH5A as E2. Dashed lines indicate positions where irrelevant lanes were cut out of the image.

**Figure S4.**
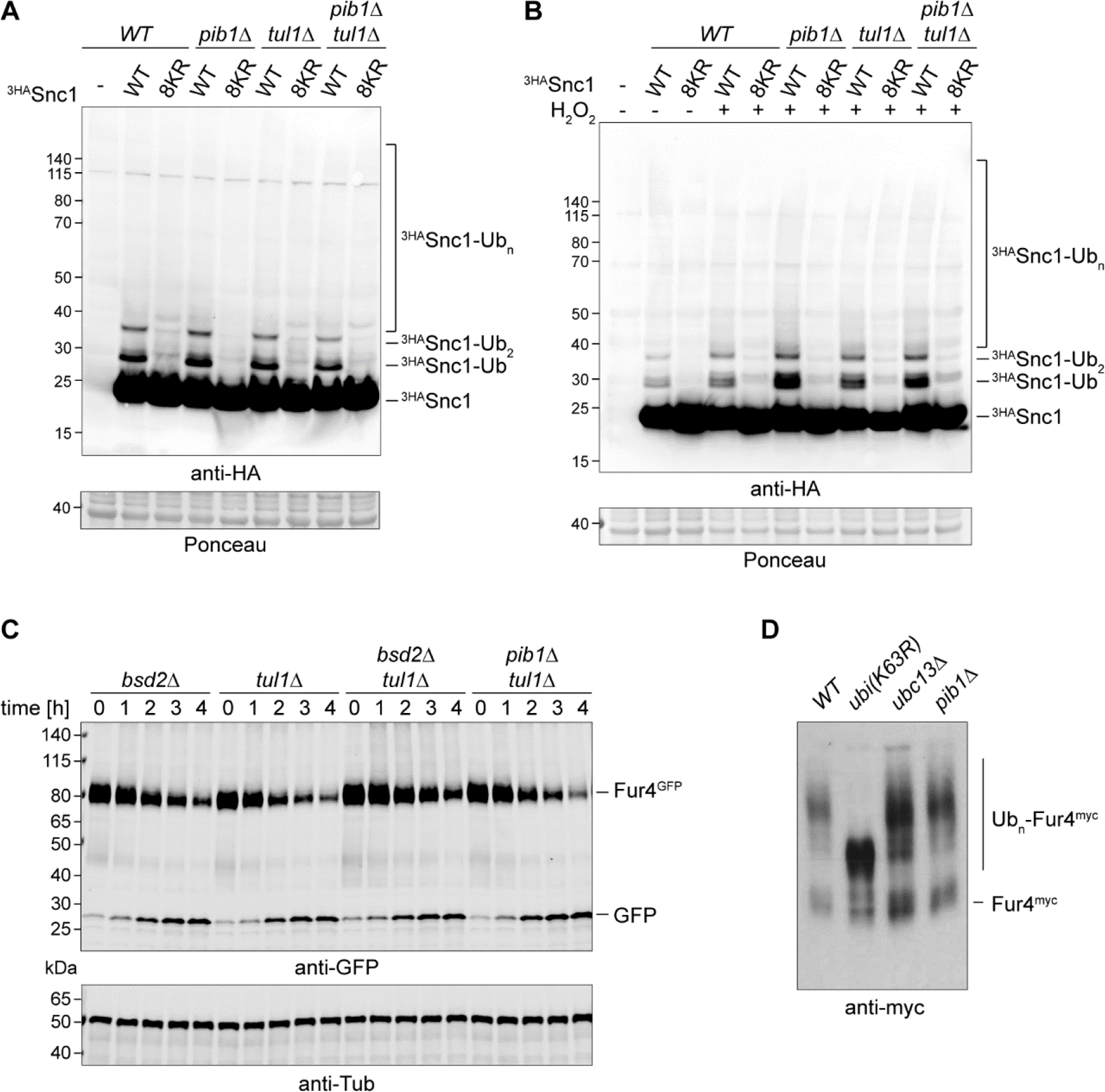
Pib1 contributes to membrane trafficking in a manner distinct from Tul1. **(A-B)** Deletion of *PIB1* and/or *TUL1* does not impair overall ubiquitylation of Snc1. Ubiquitylation of ^3HA^Snc1 (WT or a lysineless mutant, 8KR), expressed from a centromeric plasmid under control of the *TPI1* promoter, was analysed in the indicated strains by western blotting (anti-HA) of total cell extracts from exponentially growing cultures in the absence of H_2_O_2_ **(A)** or after incubation with 2.4 mM H_2_O_2_ at 30°C for 45 min **(B)**. Ponceau staining of the membrane served as loading control. **(C)** Tul1 does not contribute to vacuolar targeting of Fur4. Vacuolar delivery assays were performed in the indicated deletion strains as described in Figure 3B. **(D)** Fur4 polyubiquitylation is not dependent on Ubc13 or Pib1. Fur4^myc^ was expressed in the indicated strains from a multicopy plasmid, and the protein was detected by western blotting of a membrane-enriched lysate with an anti-myc antibody.

**Table S1.**
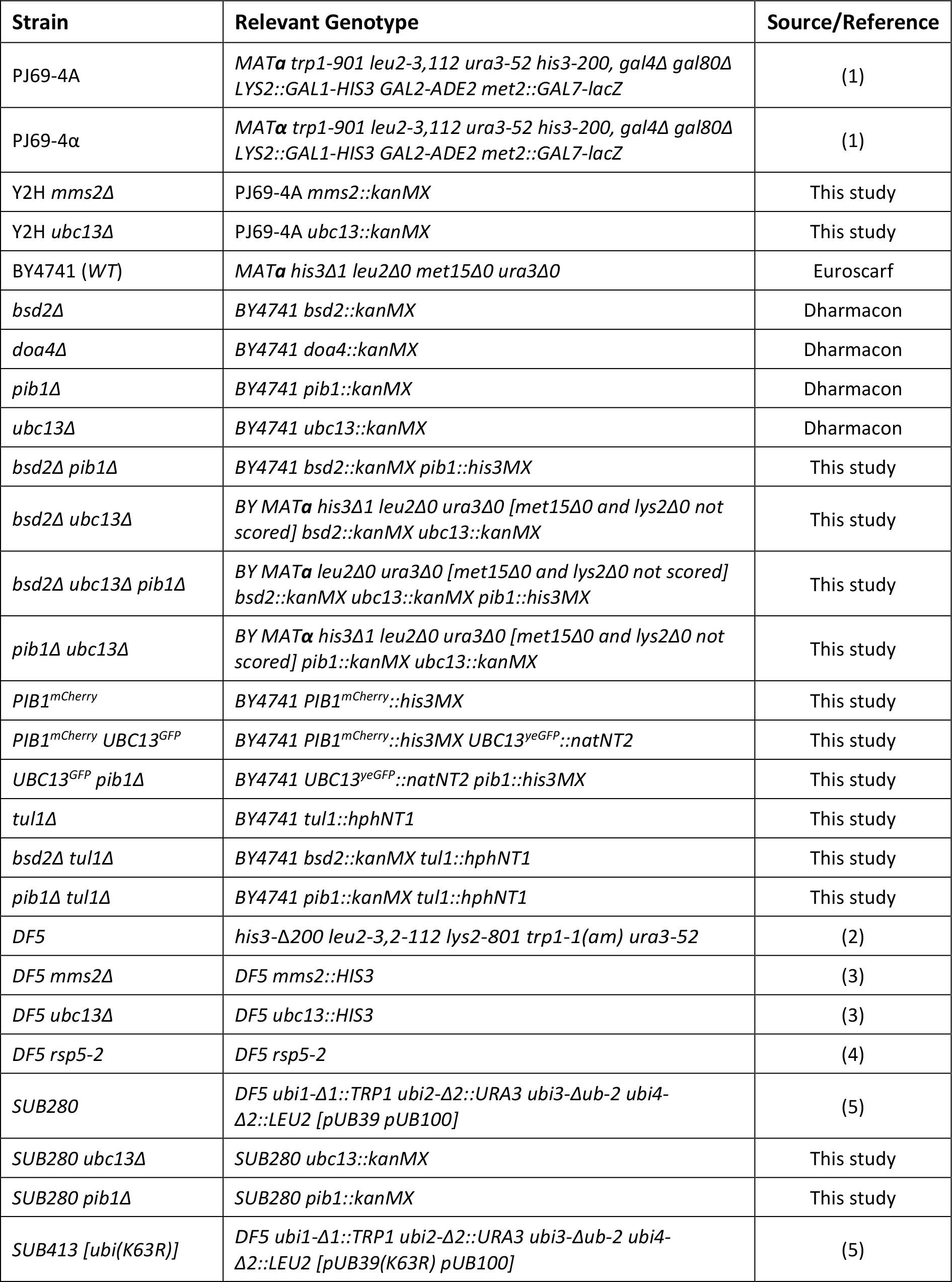
Yeast strains used in this study.

---

**Table S2.**
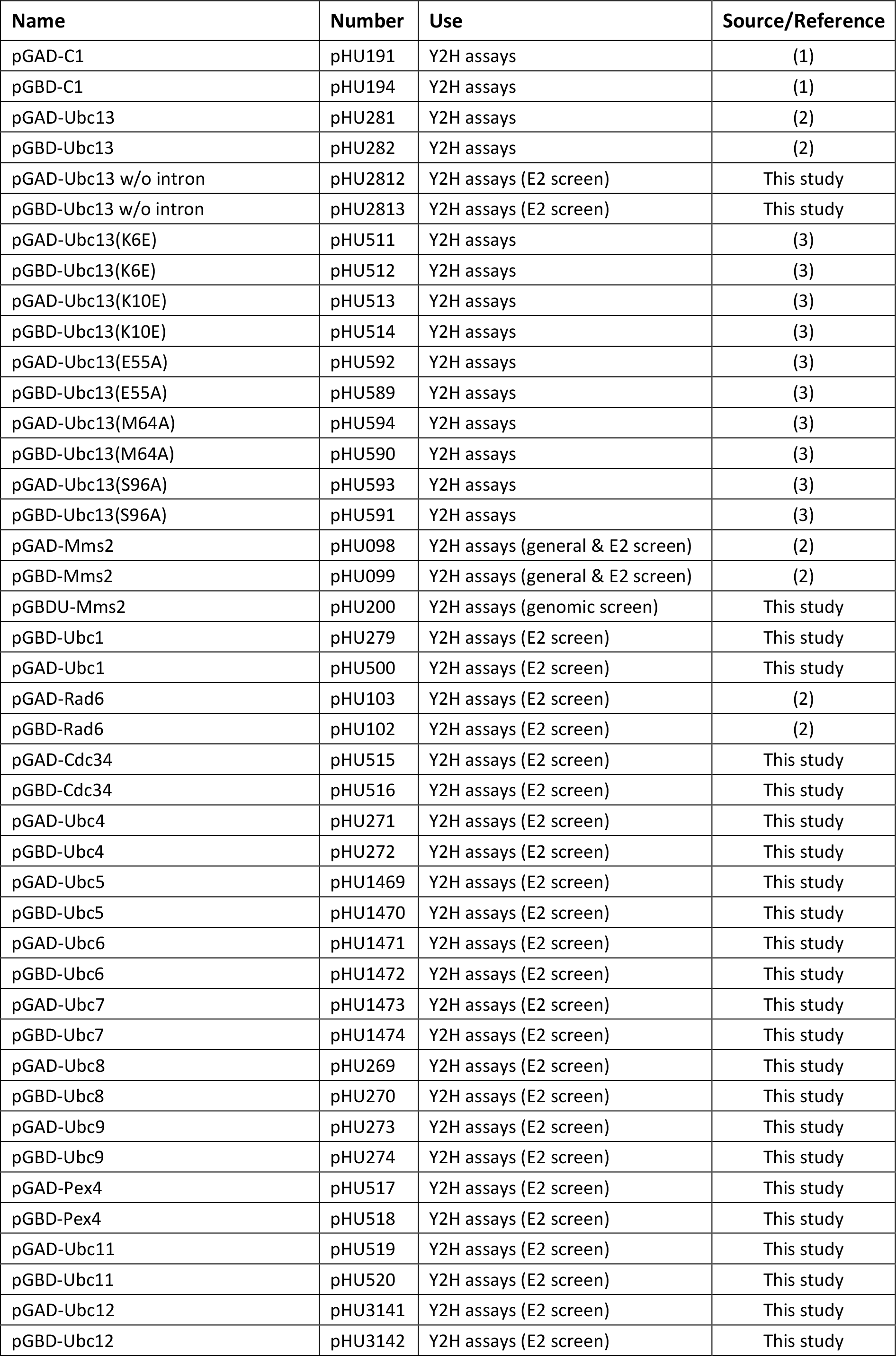
Plasmids used in this study.

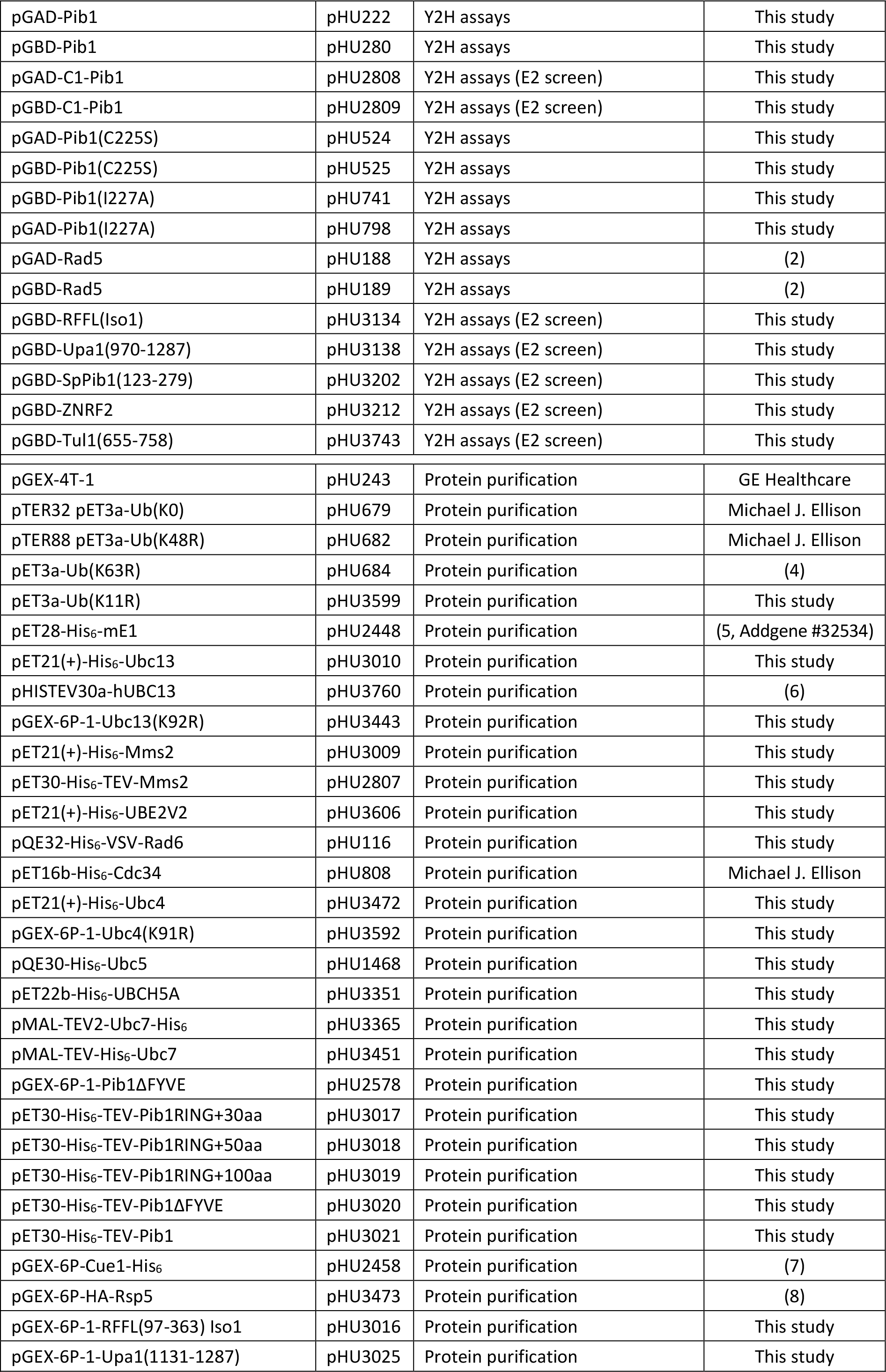

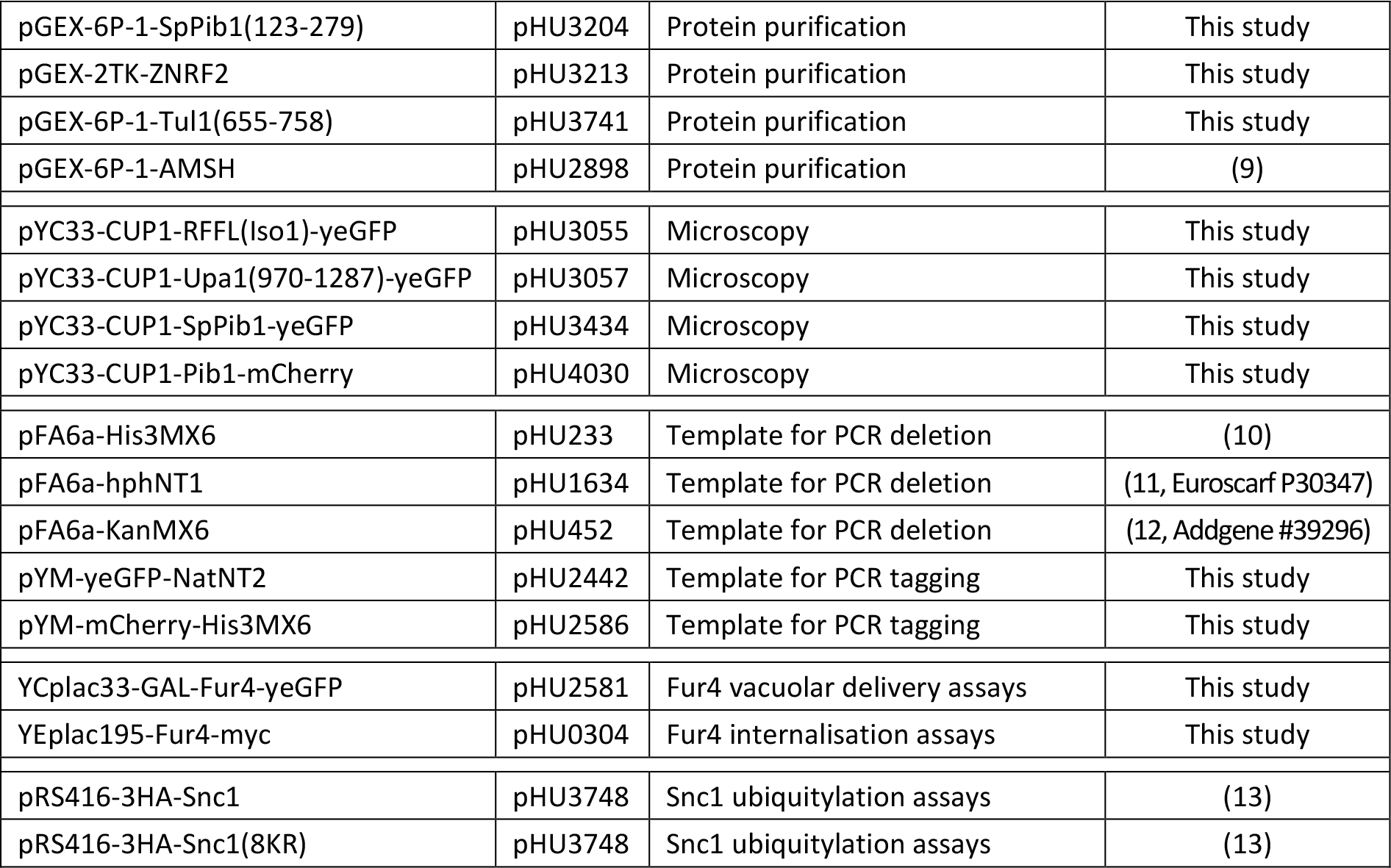

**Table S3.**
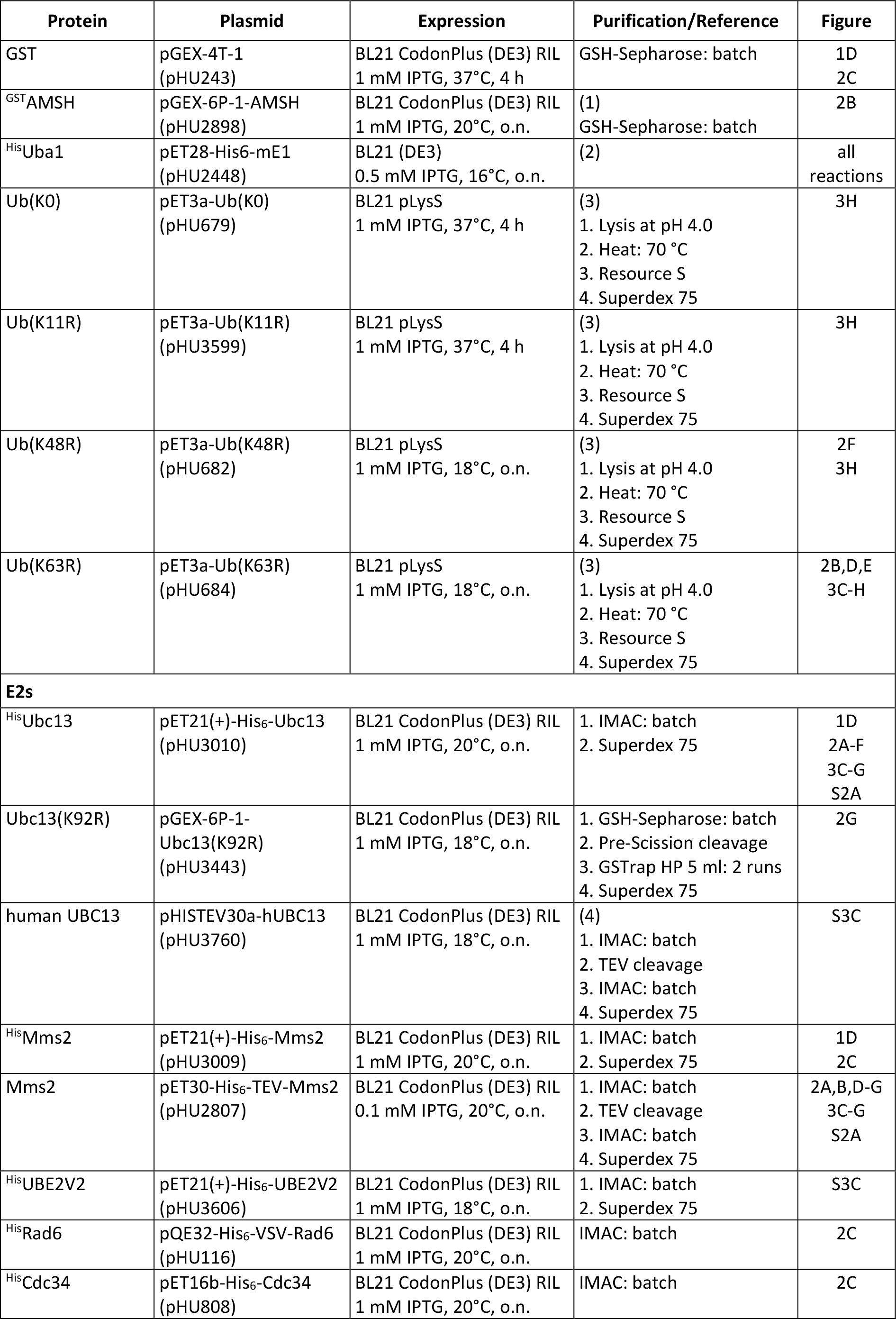
Protein purifications.

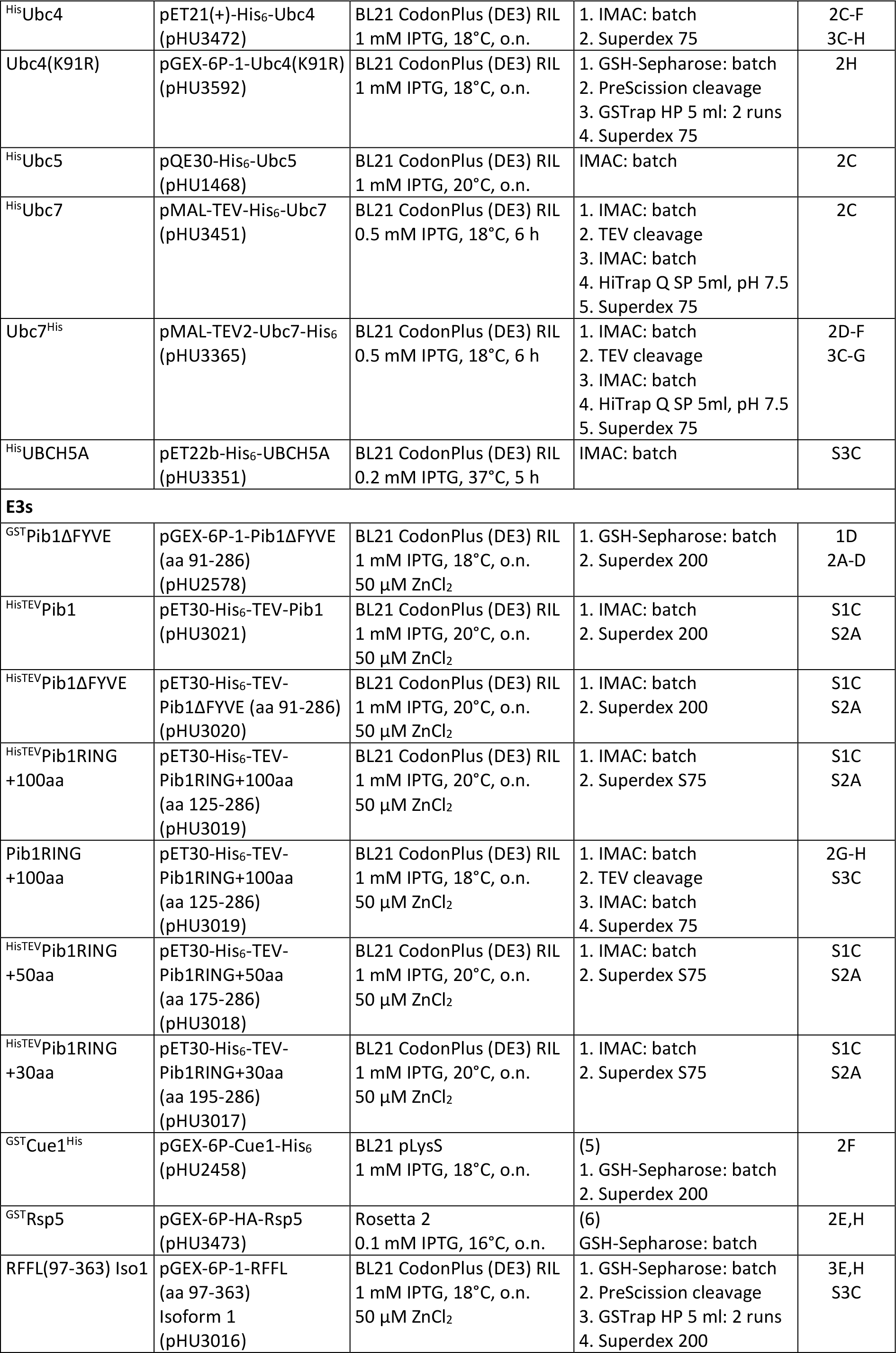

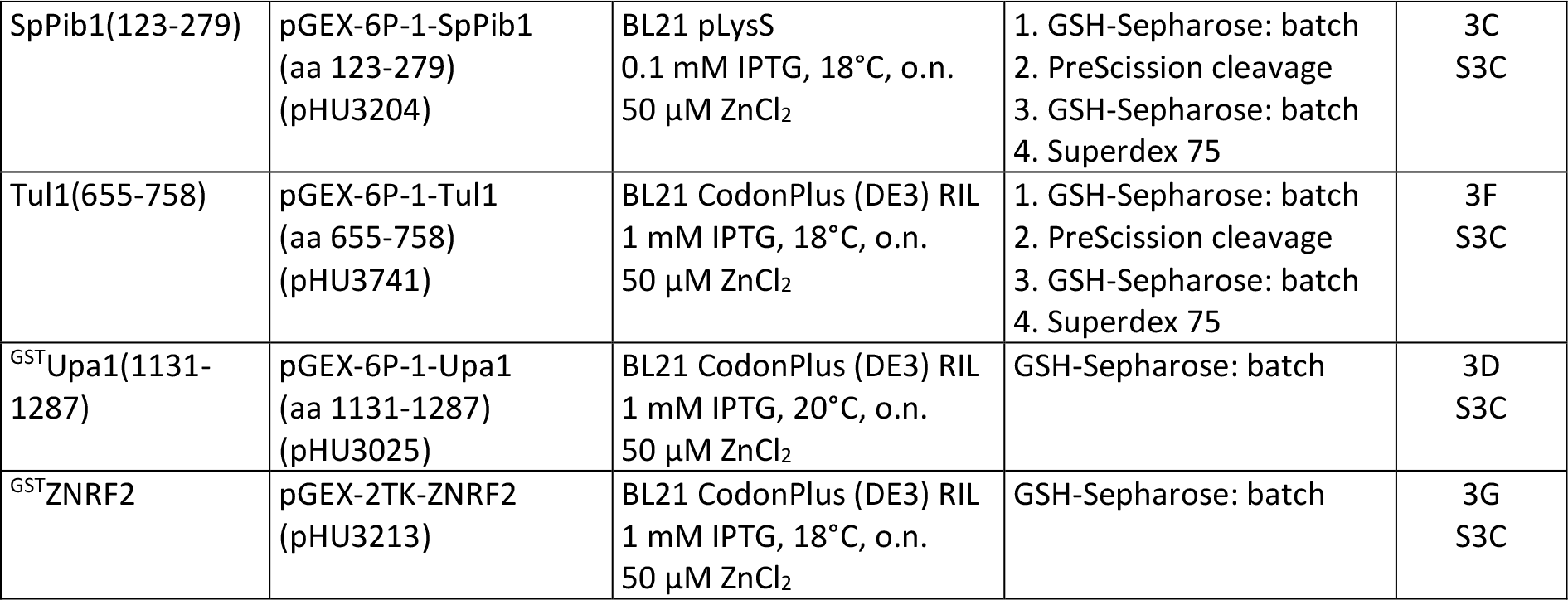

